# De novo designed ice-binding proteins from twist-constrained helices

**DOI:** 10.1101/2022.12.09.519714

**Authors:** R.J. de Haas, R.P. Tas, D. van den Broek, H. Nguyen, A. Kang, A.K. Bera, N.P. King, I. K. Voets, R. de Vries

**Author notes:** **Author Contributions:** RJdH, RPT, IKV and RdV designed the research. RJdH and RdV designed the ice-binding proteins. RJdH performed optimalization and production of recombinant proteins. RPT and DvdB performed ice-recrystallization inhibition and ice shaping assays. HN and AK screened and optimized crystallization conditions. AKB solved the crystal structure of TIP-99_a_. NPK, IKV and RdV supervised and coordinated the research. RJdH, RPT, IKV and RdV wrote the manuscript. All authors edited and accepted the manuscript. **Competing Interest Statement:** RJdH, RPT, IKV and RdV are co-inventors in a patent application (European patent application No. 22200103.4) covering twist-constrained alpha-helical ice-binding proteins described in this work. The remaining authors declare no competing interests.

## Abstract

Attaining molecular-level control over solidification processes is a crucial aspect of materials science. To control ice formation, organisms have evolved bewildering arrays of ice-binding proteins (IBPs) but these have poorly understood structure-activity relationships. We propose that reverse engineering using *de novo* computational protein design can shed light on structureactivity relationships of IBPs. We hypothesized that the model alpha-helical winter flounder antifreeze protein (*wf*AFP) uses an unusual under-twisting of its alpha-helix to align its putative ice-binding threonine residues in exactly the same direction. We test this hypothesis by designing a series of straight three-helix bundles with an ice-binding helix projecting threonines and two supporting helices constraining the twist of the ice-binding helix. We find that ice recrystallization inhibition by the designed proteins increases with the degree of designed under-twisting, thus validating our hypothesis and opening up new avenues for the computational design of ice-binding proteins.

**Significance Statement:** Ice-binding proteins (IBPs) modulate ice nucleation and growth in cold-adapted organisms so that they can survive in ice-laden environments at (sub)freezing temperatures. The functional repertoire of IBPs is diverse, ranging from inhibition of recrystallization and freezing point depression to shaping of ice crystals and ice nucleation. Precisely how these activities arise from the structure and ice-binding properties of IBPs is poorly understood. We demonstrate through *de novo* computational protein design that constraining the twist of an ice-binding helix is a key feature determining its ice-binding activity, opening new avenues for the design of synthetic IBPs with activities tailored to the requirements of specific applications, such as cell and tissue cryopreservation.

## Introduction

Control over solidification processes is a key aspect of materials science. For crystallization from the liquid state, or from solution, an important mechanism for control is via molecules that preferentially bind to certain crystal planes. Nature has evolved a wide array of molecules that exert control over crystallization processes by preferential binding to certain chemical groups in materials and to certain crystal planes. Examples include not only biomineralization, but also control over the crystallization of liquid water into ice, which is crucial for the many organisms that live at sub-zero temperatures^1,2^.

The latter is in part made possible by ice-binding proteins (IBPs) that influence growth, shaping, and nucleation of ice crystals^1,3^. Natural ice-binding proteins display highly diverse structures: many ice-binders have alpha-helical or beta-solenoid structures, but others are intrinsically disordered and heavily glycosylated^3^. The amazing structural diversity of IBPs is just one factor that has hampered the development of a general molecular-level understanding of the activities of IBPs. Another complicating factor is that ice binding affinity and facet-specificity alone are not sufficient to explain the activities of IBPs, such as freezing point depression, ice-recrystallization inhibition (IRI) and nucleation enhancement or suppression.

To test hypotheses and theories for the activities of IBPs, it would be very helpful to *de novo* design model IBPs, which may allow more straightforward elucidation of structure-activity relationships. *De novo* designed IBPs may also provide advantages for eventual technological applications: natural IBPs do not necessarily satisfy constraints imposed by technological applications such as high thermal stability and low production cost, and *de novo* design may allow consideration of these and other constraints during design.

While structure-activity relationships are still obscure for many IBPs, their molecular geometry when binding to specific ice-planes is starting to be understood, in particular through molecular simulation^4^. Such insights are a suitable starting point for the *de novo* design of IBPs. One of the best studied and simplest natural IBPs is the winter flounder AFP (*wf*AFP). It is a member of the so-called type-I IBPs. The *wf*AFP protein is a 37-residue, alanine-rich, almost straight alpha helix, which is thought to bind especially strongly to the pyramidal {2021} plane of ice, and thus shape ice into characteristic bipyramidal crystals^5,6^ (**Fig. 1A**). Helical type-I IBPs such as *wf*AFP have characteristic 11-mer consensus amino acid repeats of the general sequence TXXXAXXXAXX, where X is often alanine^7^. It is proposed that *wf*AFP binding to the {2021} plane of ice is driven by the close match between the 16.7 Å threonine spacing in the 11-mer repeat and the 16.5 Å spacing between oxygen atoms on the {2021} plane^8,9^. Threonines on *wf*AFP can be mutated to valines without much loss of activity, but mutating them to serines abolishes activity, suggesting van der Waals interactions at those positions are crucial^10^.

**Figure 1.**
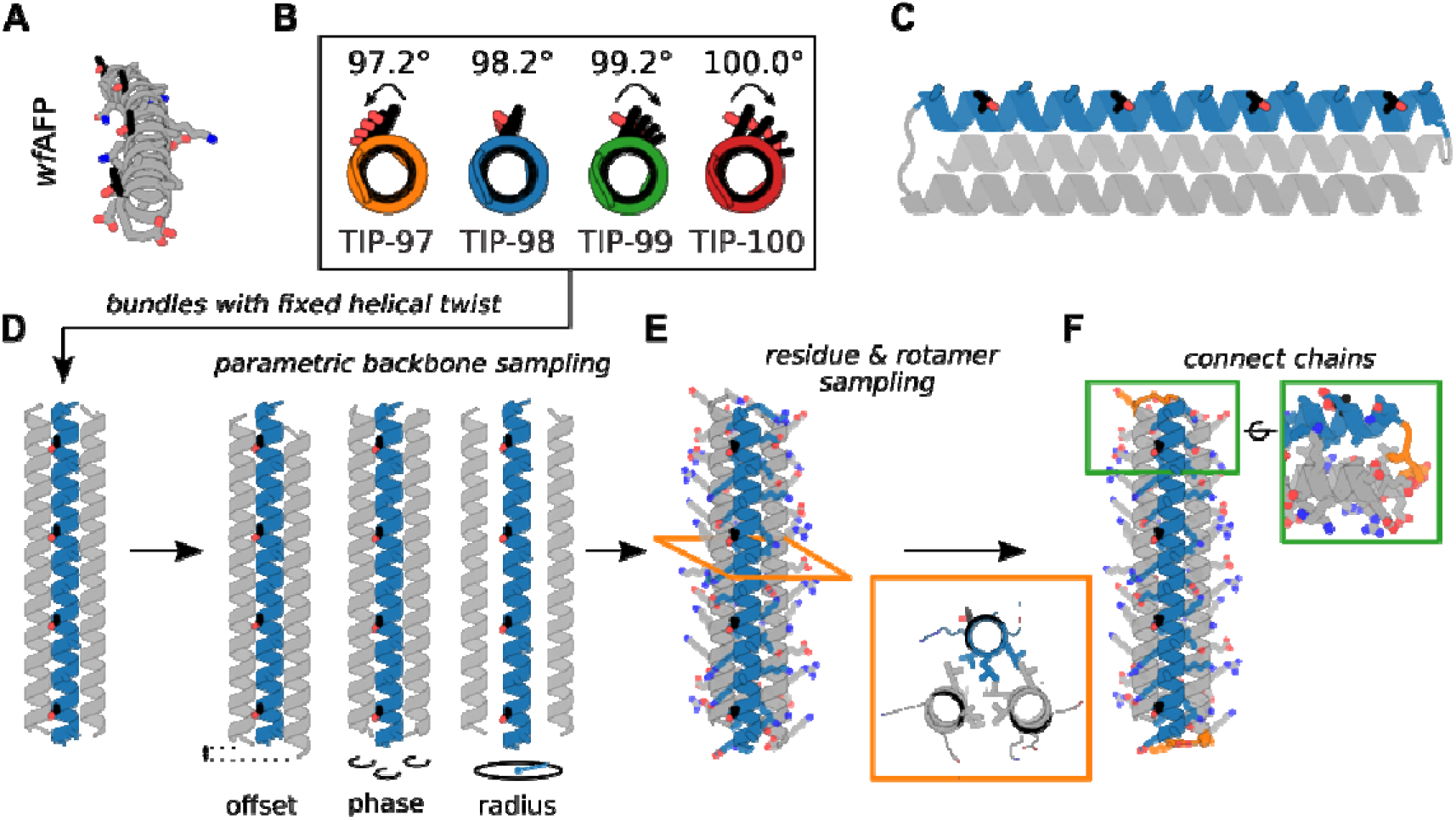
Design of <u>T</u>wist-constrained <u>I</u>ce-binding <u>P</u>roteins (TIPs). **A)** *wf*AFP crystal structure (PDB ID: 1WFA), comprising four repeats of the minimal putative ice-binding 11-mer consensus sequence TXXXAXXXAXX. The four threonine residues (black) all point in the same direction. **B)** Straight helical configurations predicted for the ice-binding helices of the designs TIP-97, TIP-98, TIP-99, and TIP-100. Arrows represent threonine (mis)alignment. **C)** TIP design concept. A helix containing a minimal ice-binding consensus sequence (threonines in black) is twist-constrained by two neighboring helices (gray). The three helices are connected into a bundle through small loops. **D)** A parametric description of helical bundles is used for backbone generation. Parameters which are sampled include offset, phase, and radius (as indicated). **E)** Fixed backbone residue and rotamer sampling is used to find optimal low-energy sequences. Insert shows an example of the tight hydrophobic core packing presumably required to properly constrain the twist. Ice-binding threonines are highlighted in black. **F)** Connection of the helices into a single bundle using 2-3 residue loops.

Previous authors have pointed out that for *wf*AFP, the threonines on the alpha-helix all point in the same direction^11^, as would be required for matching the structure of the ice {2021} planes. As far as we know however, it has not yet been pointed out that this is atypical for straight alpha helices. The ideal, straight alpha helix has a helical twist of 3.6 residues per helical turn, or 100° per residue (**Fig. S1**). For 11 residues, this implies 3.06 full rotations, or a rotation of one threonine to the next of 20°. To have exactly 3 helical rotations per 11-mer, the *wf*AFP is undertwisted, having an average helical twist of 98.2° per residue (**Fig. 1B**). Similar considerations apply to other type-I IBPs such as shorthorn sculpin AFP (*ss*AFP) (**Fig. S2**). We hypothesize this under-twisting is crucially related to their ice-binding activity. Here we take this observation as the basis for *de novo* design of a type-I model IBP.

For *wf*AFP and other type-I IBPs it is not clear how the constraint on helical twist arises from its sequence. To constrain the twist of a single putative ice-binding helix, we here propose to incorporate it in a straight helical bundle. We keep the number of helices in the bundles as low as possible. Most natural two-helix helical bundles (coiled coils) are non-straight and have a nonzero super-helical twist^12^. Therefore, we use straight three-helix bundles as scaffolds for the design of <u>T</u>wist constrained <u>I</u>ce-binding <u>P</u>roteins, or TIPs.

Relatively simple equations exist that describe the backbones of bundles of alpha-helices in terms of a few parameters, such as the number of helices, the helical twist, and a few others^13^. These equations allow for the rapid *in silico* construction of large libraries of alpha-helical backbones, which can be populated with amino-acid rotamers in a subsequent design step. Such parametric backbone generation, followed by connection of the helices with small loops and sequence design of the backbones using Rosetta, was used by Huang et al. to design highly idealized and extremely stable *de novo* helical bundles^13^. Huang et al. also designed extremely stable straight three-helix bundles, but in their designs, the twist of the helices was not constrained to a predetermined value^13^. We follow the approach of Huang et al. to design straight TIPs consisting of three-helix bundles with a twist-constrained ice-binding helix (TXXXAXXXAXX)_4_ and two supporting helices that are not ice-binding, but instead act to constrain the twist of the ice-binding helix to a designed value (**Fig. 1C**).

To test the hypothesis that helical twist is an important determinant for ice-binding, we first design a series of TIPs based on just the minimum 11-mer consensus sequence TXXXAXXXAXX, with varying helical twist of the ice-binding helix in the range of 97.2°–100° per residue. As we will demonstrate, this first series of TIPs have an ice recrystallization inhibition (IRI) activity that increases with the degree of designed under-twisting. Next, for the optimal twist, we investigate the impact of judiciously replacing surface exposed ‘X’ residues in the minimal consensus sequence with alanines. This further increases IRI activity of the TIPs, to a level comparable to that of the *wf*AFP on which the *de novo* designs were based, thus providing a first example of a nature-inspired *de novo* designed ice-binding protein.

## Results

### TIPs with minimal ice-binding consensus motif

In our first series of TIPs we chose to use the minimal consensus motif TXXXAXXXAXX, without constraining the identities of any of the ‘X’ residues. We reason this will yield proteins with favorable solution properties, because alanine-rich protein surfaces can increase aggregation propensity via hydrophobic interactions^14^. We chose four target twist angles for the ice-binding helix. The first was the ideal twist angle 98.2° that yields exactly three helical rotations for an 11-mer motif. Next we chose twist angles that are, respectively, 1° higher and lower than this. Finally we also chose the twist angle of a typical straight alpha helix of 100°. The corresponding proteins were designated TIP-97, TIP-98, TIP-99 and TIP-100.

Helices in helical bundles are characterized by a major (super-helical) twist ω_0_ and a minor (alpha-helical) twist ω_1_. We intended to design straight helical bundles, hence we fixed the target value of the major helical twist for all helices to ω_0_ = 0° and fixed the target values for the minor helical twist for all helices to ω_1_ = 97.2°, 98.2°, 99.2° and 100° for TIP-97, TIP-98, TIP-99, and TIP-100 respectively. Note that even the 1° change of the twist angle away from the optimal value of 98.2° already leads to an accumulated angle between the first and last threonine of 33.6° (**Fig. 1B**), which we hypothesized would lead to less precise matching to the ice-lattice plane, and hence to lower binding affinities for these ice planes.

We followed the general approach developed by Huang et al. for parametric design of helical bundle backbones, coupled to Rosetta Design for populating the backbones with aminoacid rotamers^13^. We sampled helical parameters for each of the three helices in the bundle independently. In total there are six parameters describing each three-helix bundle: offset (2), phase (3), and the bundle radius (**Fig. 1D**). The ranges of parameter values from which we sampled are given and motivated in **Supplementary Information.** Through combinatorial sampling we generated large sets of 15,000–30,000 backbones per twist. The large set of very diverse backbones is crucially important for finding a backbone that allows for a well-packed bundle core that properly constrains the ice-binding helix. Next we generated atomistic models by populating each backbone with amino acid rotamers using Rosetta fixed backbone design (details in **Supplementary Information**). The identifies of all amino acids were left free in the design procedure, except for the threonines and alanines in the tandem repeat of the consensus 11-mer motif TXXXAXXXAXX on the central ice-binding helix.

Models were scored using the Rosetta REF2015 energy function^15^ and subjected to a packstat filter^16^ that removed designs with poor core packing, since these likely would not be able to properly constrain the twist of the ice-binding helix to its target value. Next, for each target value of the twist, we selected the 1% of models with the lowest energy score (150–200 models per target twist value). RosettaRemodel^17^ was then used to design short loops connecting the helices in the bundle using the KIC^18^ protocol (details in **Supplementary Information**). Per designed and selected backbone, 200 loop closure attempts were made. The 1% of chain-connected models with the lowest energy score (75–100 models) were selected for further consideration.

Selected chain-connected models were relaxed using the Rosetta Relax application (with flexible backbone)^19,20^. To assess how well the designed sequence specifies the target structure, we calculated the Cα-RMSD between the designed and relaxed structures of each model. This was used to further narrow down the selection of models: from the 75–100 chain-connected models per target twist value, we selected the models with Cα-RMSD < 1.5 Å, and from these we selected 10-15 models having the lowest energy score.

These models were next folded from the extended chain using the Rosetta Abinitio Relax^21^ application to probe the folding energy landscape (details in **Supplementary Information**). Designs with folding funnels towards a single low-energy state <2.5 Å Cα-RMSD from the designed model were selected for experimental characterization (**Fig. S3**): four TIP-100 designs, four TIP-99 designs, seven TIP-98 designs, and four TIP-97 designs.

We found excellent agreement between the helical twisting of the ice-binding helix of the designed and relaxed models for these designs (**Fig. S4A**), indicating that at least within the framework of the Rosetta REF2015 energy function, we managed to design sequences that adequately constrain the twist of the ice-binding helix. As an additional check, we also folded the designed sequences using AlphaFold2 and found excellent agreement for the twist of the ice-binding helix as compared with the Rosetta ab initio predictions (**Fig. S4B**) as well as mostly good agreement in the overall predicted structures. Notable differences were found for the TIP-98 models, where AlphaFold2 predicted a different topology of the helical bundles, and for the TIP-97 models, where AlphaFold2 predicts (with low confidence) a nonzero major helical curvature (**Fig. S5 and Table S2**).

Next we proceeded to experimental characterization. For TIP-100, 2/4 designs expressed well and were soluble. However, these designs were not monomeric as assayed by size exclusion chromatography (SEC), and were not considered further. For TIP-99, 4/4 designs expressed well and were soluble. Of these, 2/4 were found to be monomeric as assayed by SEC. Both designs, referred to as TIP-99_a_ and TIP-99_b_, were alpha-helical by circular dichroism (CD) and stable up to at least 95°C (**Fig. 2A-B**). For TIP-98, 2/7 designs expressed well and were soluble. One of these was monomeric as assayed by SEC. This design was also found to be alpha-helical by CD and stable up to at least 95°C (**Fig. 2C**). Finally, for TIP-97 4/4 designs were soluble, and two of these were monomeric by SEC. One of these designs was tested by CD and found to be alpha-helical and stable until ~85°C. As opposed to the other tested designs, CD analysis of this protein indicated incomplete re-folding upon cooling (**Fig. 2D**).

**Figure 2.**
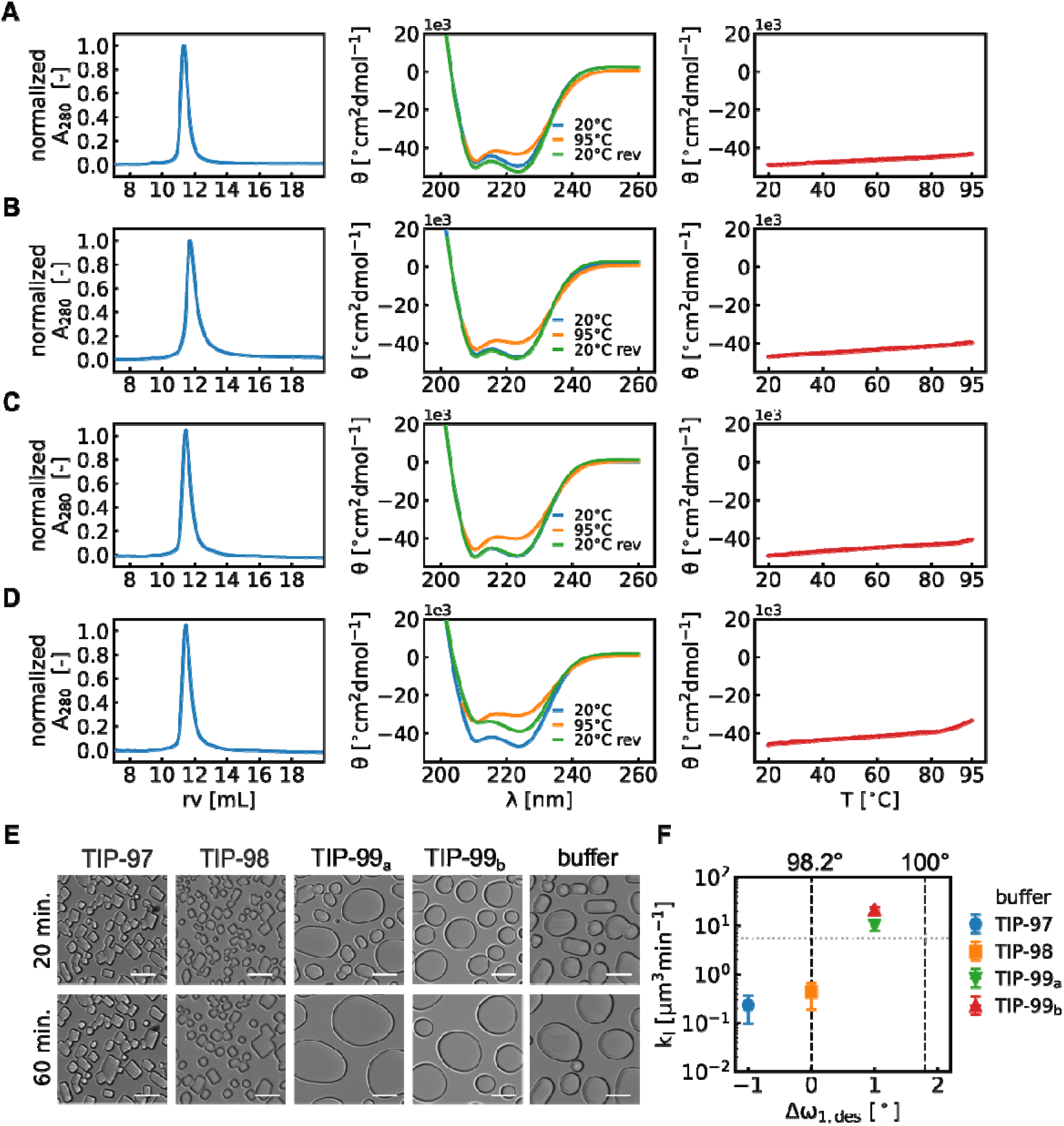
Experimental characterization and activity TIP proteins. Biochemical characterization of **A)** TIP-97, **B)** TIP-98, **C)** TIP-99_a_, **D)** TIP-99_b_. Left: size-exclusion chromatography on a Superdex 75 10/300 column in PBS+ (10 mM phosphate pH 7.4 + 300 mM NaCl), demonstrating proteins elute at the expected retention volume (rv) of ~12 mL, corresponding to a 17 kDa globular protein. Middle: circular dichroism spectra. Molar ellipticity θ) versus wavelength (λ) at 20°C, 95°C and 20°C reversed (20°C rev) demonstrating alpha-helical secondary structure and high thermal stability. Right: molar ellipticity (θ) at 222 nm during temperature ramp from 20°C to 95°C (1°C/min). **E)** Microscopy image of ice crystals at 20 min. (top) and after 60 min. (bottom) exposure to −7°C in the absence (buffer) or presence of TIPs at 60 μM, demonstrating ice-recrystallization inhibition (IRI) for TIP-97 and TIP-98. Scale bar: 20 μm. **F)** Quantitative IRI analysis. Speed of growth of ice-crystals [μm^3^ min^-1^] during Ostwald ripening at a fixed protein concentration of 60 μM as a function of the deviation from proposed ideal twist angle.

We analyzed whether TIP-97, TIP-98, TIP-99_a_, and TIP-99_b_ (**Table S1**) could inhibit the recrystallization of small ice-crystals (IRI activity). In this assay, a dispersion of small ice-crystals is first created, and the growth of these crystals at −7°C is observed under a microscope. Typically, a first phase with mainly rapid coalescence is followed by a second slower phase of mainly Ostwald ripening. Potential IRI activity of the proteins is manifested in the second phase as a reduced speed of growth of the ice-crystals. Example microscopy images in **Fig. 2E**. very clearly demonstrate that TIP-97 and TIP-98 proteins significantly retard the Ostwald ripening of the ice crystals. Next, to analyze how IRI activity depends on the (designed) degree of twisting of the ice-binding helix, image analysis was performed to obtain the speed of growth of the ice-crystals at a fixed protein concentration of 60 μM. As shown in **Fig. 2F**, we find that IRI activity increases with the degree of designed under-twisting of the ice-binding helix, with the TIP-99 proteins exhibiting lower activity than the TIP-98 and TIP-97 proteins. According to our hypothesis, the largest IRI activity would have been predicted for TIP-98. This expectation is not entirely borne out by the experiments, in the sense that we do not observe the IRI activity decrease for (designed) degrees of under-twisting beyond the supposedly optimal value of 98.2° per residue. We speculate that this may either be caused by small structural differences between experimental and target designed structures, or by other potential factors that contribute to ice-binding, apart from the degree of under-twisting of the ice-binding helix.

To better understand the structural basis of the IRI activity, we set out to crystallize our TIP constructs. Extensive screening resulted in high quality diffraction of TIP-99_a_ for which a structure was resolved at a resolution of 2.3 Å (**Fig. 3**). The crystal structure shows that TIP-99_a_ maintains the designed helical twisting of 99.2° per residue and closely matches the Rosetta-relaxed model, with a backbone RMSD of 1.12 Å. For the ice-binding helix, the deviation is slightly smaller (backbone RMSD of 0.67 Å). The high degree of agreement between the experimental and designed structures provides confidence in our design protocol for these unusual twist-constrained alpha-helices.

**Figure 3.**
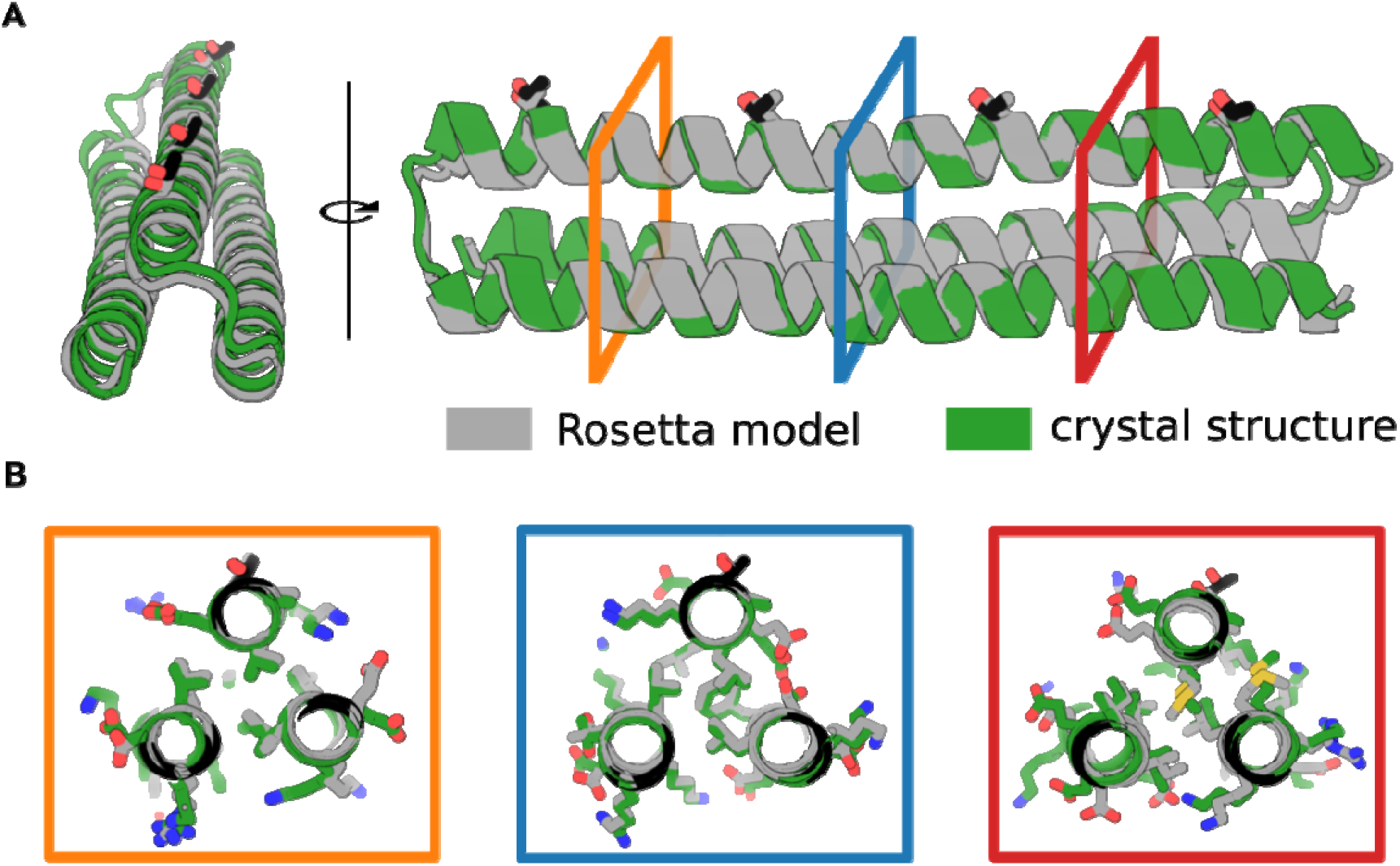
Crystal structure of TIP-99_a_. **A)** Cartoon diagram showing TIP-99_a_ crystal structure (green) and Rosetta-relaxed design model (gray) with an overall backbone RMSD of 1.12 Å across 405 aligned atoms. The crystal structure of TIP-99_a_ (PDB ID: 8EK4) was solved to a resolution of 2.3Å. The helical twist of the ice-binding helix and the ice-binding residues are maintained at 99.2° per residue with a 0.67Å backbone RMSD of the ice-binding helix between 132 aligned atoms in the design model and crystal structure. **B)** Three views at different locations in the helical bundle showing rotamer packing in the crystal structure, particularly in the core, closely matches the design model. The ice-binding threonines are at the top of each image are displayed as sticks, with threonines of the crystal structure highlighted in black and threonines of the Rosetta-relaxed design model in gray.

### Improving ice-binding activity by expanding the ice-binding motif

Having found that the IRI-active TIP-98 protein was highly thermostable and soluble, we reasoned there should now be enough design freedom to constrain the twist of the ice-binding helix while at the same time optimizing the nature of the amino acids in the vicinity of the putative ice-water interface to further enhance IRI activity. The ‘X’ residues in the minimal type-I IBP consensus sequence of TXXXAXXXAXX are often alanine: a pattern search for the consensus 11-mer motif for the type-I IBPs wfAFP and the ssAFP (PDB ID: 1Y03) yields a dominant motif of TAAAAAAAAAAA (See logo plot in **Fig. S6**). Indeed, we expect that mutating the high-entropy surface rotamers preferably designed by Rosetta (lysines, arginines, glutamic acids, etc.) to small, hydrophobic alanines, may help to better stabilize water molecules surrounding the ice binding plane on the protein.

In the TIP-98 protein, ‘X’ residues at positions 2 and 8 of the 11-mer motif are expected to be both solvent-exposed and in close contact with the ice-water interface as they are closest in distance and orientation to the ice-binding threonines. In an attempt to improve the IRI activity of TIP-98, we therefore created the mutants TIP-98^2A^ and TIP-98^8A^ with alanines at, respectively, positions 2 and 8 in the type-I 11-mer repeat motif, as well as the double mutant TIP-98^2A8A^ (**Fig. 4**). Using Rosetta Design, model structures were obtained for TIP-98^2A^, TIP-98^8A^ and TIP-98^2A8A^. We found that for all of these, the ice-binding helix retained the major helical twist of ω_0_ = 0° and minor helical twist of ω_1_ = 98.2° thought to be crucial for ice-binding (**Fig. S7D**). All three alanine mutants expressed as soluble, monomeric, alpha-helical proteins, with thermal stability up to at least 95°C (**Fig. S7A-C**).

**Figure 4.**
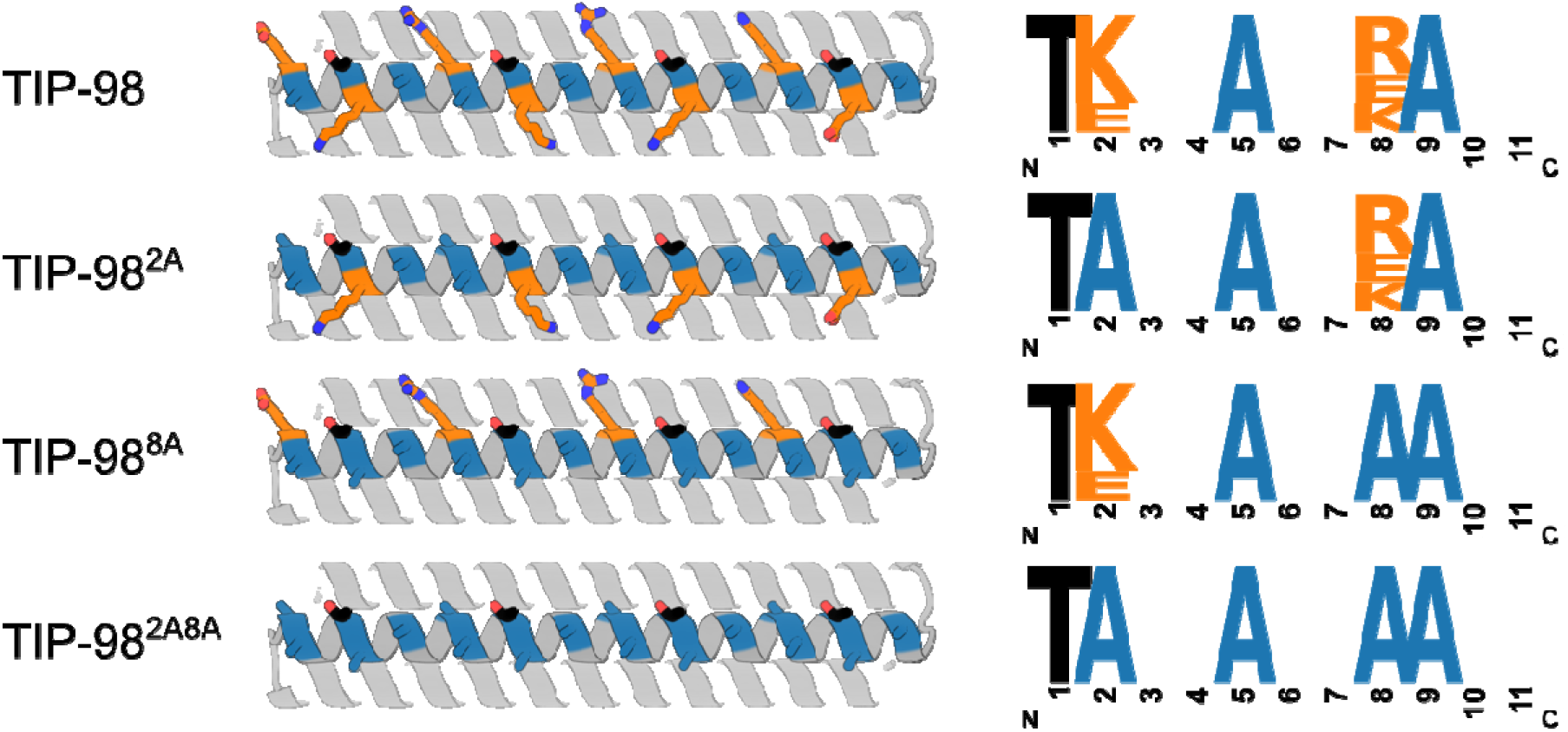
Expanding the minimal consensus sequence to increase activity. Predicted structures and ice-binding sequence motifs for TIP-98, TIP-98^2A^, TIP-98^8A^, and TIP-98^2A8A^. The amino acids at positions 2 and 8 (orange) were mutated to alanine (blue) individually and in combination to improve ice binding. Ice-binding threonines are highlighted in black.

For this second series of TIP proteins, a more quantitative characterization of IRI activity was performed. For all proteins, the speed of growth of the ice-crystals during Ostwald ripening was measured as a function of protein concentration in order to more precisely determine a critical concentration required for IRI activity. Example microscopy images at fixed TIP concentration of 20 μM are shown in **Fig. 5A** and quantitative IRI results at various TIP concentrations are summarized in **Fig. 5B.** We found that the original TIP-98 only has a quasi-critical, rather than a true critical concentration: at high TIP-98 concentrations, the speed of growth of the ice-crystals did not reduce to zero but instead attained a low, non-zero value. The value of this quasi-critical concentration is approximately 30 μM. In contrast, the alanine mutants have a true critical concentration, beyond which growth of the ice-crystals is fully arrested. This true critical concentration for IRI was highly similar for the different alanine mutants and is approximately 10 μM.

**Figure 5.**
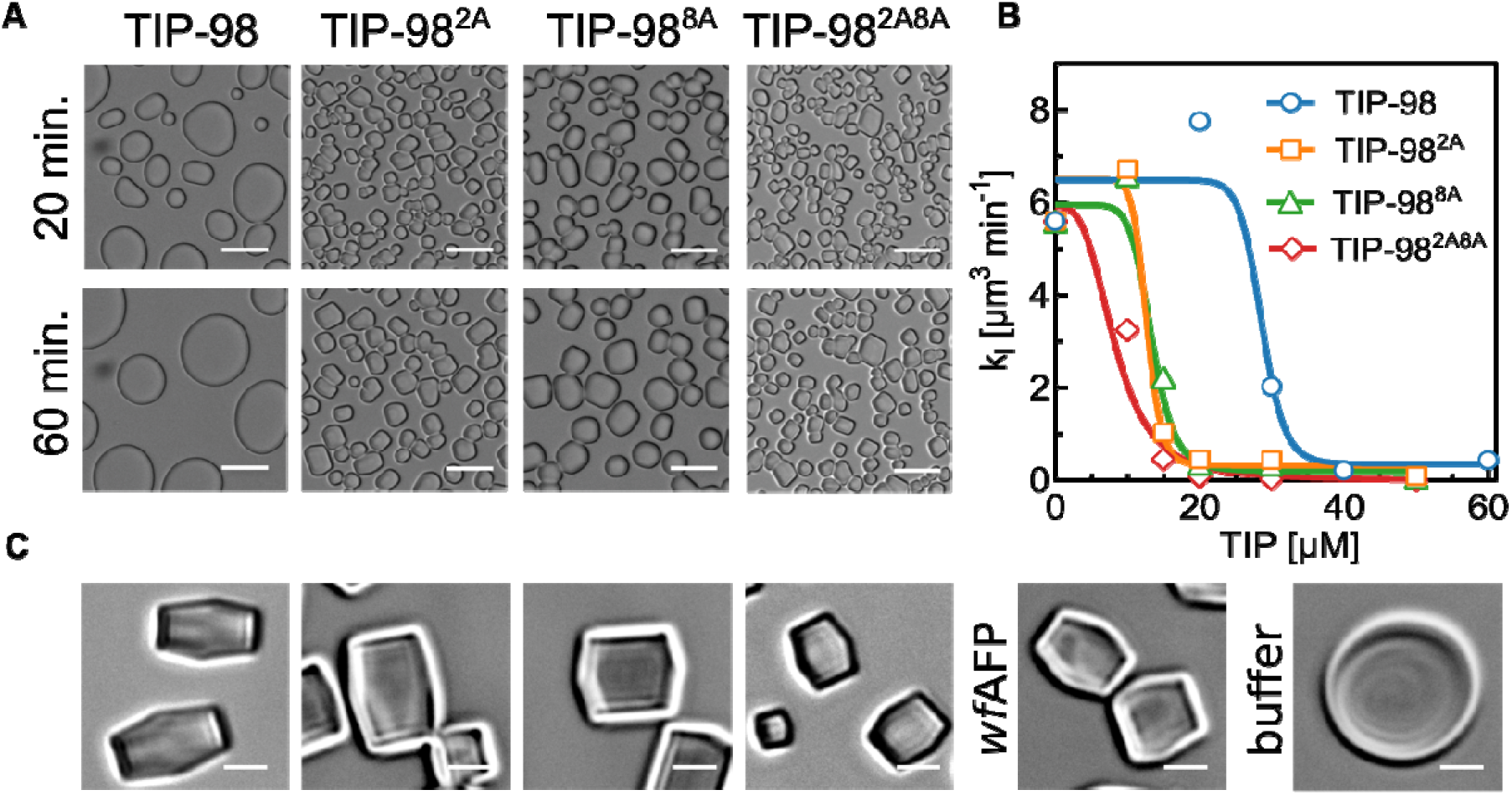
Improved activity with “maximal” consensus sequence. **A)** Ice Recrystallization Inhibition (IRI) microscopy images at 20 μM. Scale bar: 20 μm. **B)** Quantitative IRI results: average speed of crystal growth rate [μm^3^ min^-1^] as function of TIP concentration. **C)** Ice-shaping into blunt-end crystals for TIP-98, TIP-98^2A^, TIP-98^8A^, TIP-98^2A8A^, and *wf*AFP at 50 μM. Absence of IBPs (buffer) leads to ice crystals with various (round) morphologies. Scale bar: 5 μm.

Finally we aimed to gain insight in the ice-planes to which the TIP-98 proteins preferentially bind. Here, crystal shaping of our designed TIP-98 variants was compared alongside that of type-I *wf*AFP that binds the pyramidal {2021} plane of ice^5,6^. These Ice-shaping experiments were performed by undercooling ice crystals at 0.2°C/min in the presence of 50 μM TIP-98, TIP-98^2A^, TIP-98^8A^, TIP-98^2A8A^ or *wf*AFP, which forced the crystals to shape. Microscopy images obtained after one minute of undercooling clearly demonstrated that TIP-98 variants induce similar (blunt) bipyramidal shaping of ice-crystals as *wf*AFP, strongly suggesting they likewise preferentially bind to the pyramidal {2021} plane of ice (**Fig. 5C**).

## Discussion

Comprehensive studies on systematic series of novel compounds^23–26^ as well as detailed molecular dynamics simulations^4,27^ have significantly advanced our understanding of the relation between atomic-level structural features and functional activity of ice-binding compounds. Yet, so far, rational design has not led to new ice-binding compounds with activities that compete with many natural ice-binding compounds, in particular IBPs.

One of the complicating factors is that for most compounds it is not possible to independently modulate composition and structural features, such as flexibility, twist, curvature, and bulkiness. As a consequence, structure-activity relations remain opaque and controlling activity through design is difficult. Our study demonstrates how computational protein design methods can be leveraged to precisely decouple protein structure from protein surface chemistry, how this leads to atomic-level insights into structure-activity relationships for natural IBPs such as wfAFP, and how these insights lead to controllable design of ice-binding activity.

Inspired by the under-twisted alpha-helical type-I AFP from winter flounder, we designed *de novo* a series of twist-constrained proteins (TIPs). We found that precisely controlling the spacing and orientation of solvent-exposed threonines on twist-constrained alpha-helices led to potent *de novo* IBPs. As hypothesized, the IRI activity of the TIPs increased with the degree of designed under-twisting, with TIPs bearing alanine residues at positions 2 and 8 (TIP-98^2A^, TIP-98^8A^ and TIP-98^2A8A^) exhibiting levels of activity comparable to that of wfAFP.

The work described here can be extended in many directions, a few of which we wish to point out here. At the most practical level, given their high thermostability and ease of recombinant production, the TIPs may be interesting scaffolds for emerging applications of IBPs in cryopreservation of cell cultures, organoids, tissues and organs^28^. Key activities sought in this field are, among others, preventing recrystallization and explosive ice growth, control of ice nucleation, and the induction of blunt crystal shapes^29^.

Here we have focused on just a single natural IBP, the alpha-helical *wf*AFP, one of the best studied representatives of the type-I IBPs. We have no doubt that, as we have discovered here, for other classes of IBPs, it is possible to uncover similar structural features that dictate activity and which can be implemented on suitably chosen protein scaffolds.

In the longer run, as we improve our mechanistic understanding of the ice-binding activities of *de novo* designed proteins, it may also become possible to design IBPs without resorting to natural templates, with possibly new types of activities that exceed those of natural IBPs. Needless to say, ice-crystal planes are extremely repetitive at the atomic scale, and therefore it is only natural to consider repeat proteins as scaffolds for designing such IBPs. For example, repeat proteins have already been designed that are lattice-matched to crystal facets in two dimensions^30^. This may also eventually lead to *de novo* design of IBPs that promote ice-nucleation, since this is thought to require very strong binding of large macromolecular objects to relatively large areas of ice crystal planes^31^.

Ambitious protein design efforts such as the design of IBPs with no resort to natural templates seem now more possible than ever, given advances made possible by the use of new deep-learning techniques such as RoseTTaFold^32^ and AlphaFold2^22^ for protein structure prediction and deep network hallucination^33,34^ and proteinMPNN^35^ for computational protein design.

In conclusion, using the type-I IBP wfAFP as a template, we have leveraged computational protein design to decouple protein structure from protein surface chemistry for IBPs, opening up the field of *de novo* design of proteins with ice-binding activity. Given the richness of natural IBP templates available, and the current promise of improved deep learningbased protein design, we expect to see rapid progress in the design and application of designed ice-binding proteins in the near future.

## Materials and Methods

### Synthetic gene construction

Gene fragments encoding TIP proteins were codon optimized using Codon Harmony (1.0.0) and obtained as synthetic DNA fragments from Twist Bioscience. The gene fragments were cloned into a modified pET-24(+) expression vector using standard restriction cloning with BamHI and XhoI restriction endonucleases. The cloned plasmids were sequence verified using Sanger sequencing and transformed into T7-Express *Escherichia coli* (NEB) via heat-shock.

### Bacterial protein expression and purification

A 25 mL terrific broth culture supplemented with 50 mg/L kanamycin antibiotic was inoculated. The starter culture is grown overnight at 37°C in a shaker and used to inoculate 1L of luberia broth (10g tryptone, 10g NaCl and 5g yeast extract) supplemented with 50 mg/L kanamycin. The culture was incubated shaking until 0.6 < OD_600_ < 0.8 at 37°C in a 2L baffled Erlenmeyer. Protein expression was induced by 1 mM isopropyl B-D-thiogalactoside (IPTG) and expression was continued at 18°C overnight. The cell broth was centrifuged at 6,000 x g and the cell pellet was resuspended in ice-cold 30 mL lysis-wash buffer (50 mM Tris-HCl pH 8.0, 300 mM NaCl, 30 mM imidazole) supplemented with 1 mM phenylmethylsulfyl fluoride (PMSF) serine protease inhibitor and a pinch of DNAseI. Cells were then lysed by sonication on ice for 7 min. with a 2s on-off duty cycle at 85% amplitude using a Qsonica Q125 with a CL-18 probe. The lysate was centrifuged at 30,000 x g for 30 min. at 4°C and the clarified supernatant was applied two times on lysis-wash buffer equilibrated Ni-NTA resin with a column volume (CV) of ~2 mL. The resin was washed with 25 CVs lysis-wash buffer and eluted with 3 CVs elution buffer (25 mM Tris-HCl pH 8.0, 300 mM NaCl, 300 mM imidazole). Eluted protein was dialyzed to PBS+ (10 mM phosphate pH 7.4 + 300 mM NaCl) and further purified by size-exclusion chromatography on a Superdex 75 10/300 (GE Healthcare) in PBS+ on an Agilent infinity II system. Protein purity was analyzed by SDS-PAGE and purified protein was concentrated to ~20 mg/mL and stored at 4°C.

### Circular dichroism

TIP proteins were diluted to 0.15 mg/mL in PBS+ in a quartz cuvette with a 1 mm pathlength. On a JASCO J-715 (JASCO Corporation) spectral scans were averaged over 20 measurements with scan rate of 50 nm/min and a response time of 2s. A spectral scan was performed at 20°C, followed by thermal ramp to 95°C at 220 nm with rate of 1°C/min. After reaching 95°C the temperature a spectral scan was performed and temperature was reversed to 20°C (20°C rev), and another spectral scan was performed. Data where the high tension was above 600V is not shown.

### Ice recrystallization inhibition

Samples were prepared by dilution of the protein to the indicated concentration in 20 wt% sucrose in PBS+. Subsequently, a 2 μL sample was applied on a 22×22 μm coverslip and a second coverslip was lowered on top of the drop so that the sample was sandwiched between the coverslips and transferred to a Nikon ECLIPSE Ci-Pol Optical Microscope equipped with a Nikon L Plan 20x (NA 0.45) objective and the Linkam LTS420 stage. This stage was controlled by the Linksys32 software. To measure the ice-recrystallization rates, the sample was first completely frozen by reducing the temperature to −40°C with 20 °C/min. After freezing the temperature was gradually increased to −10°C with 10 °C/min and then further to −7°C with 1 °C/min upon which individual crystals could be observed and the sample was stabilized. Recrystallization was monitored by obtaining an image each minute. IRI rates were then analyzed using ImageJ and Matlab. In brief, the 8-bit images of the ice-crystals were subjected to the bandpass filter, enhance contrast and subtract background function of imageJ. Subsequently, the bright signal at the edges and then the individual crystals were isolated by the autoThreshold and Convert to Mask function. Analyze Particles was used to obtain the area of each crystal. This data was imported into Matlab upon which the radius was calculated in order to determine the corresponding spherical volume of each crystal. Recrystallization growth rates were determined by applying a linear fit to the resulting ice-volume as a function of time traces.

### Ice shaping assay

To monitor ice-shaping in the presence of the various TIPs, the sample was prepared as for the IRI assays but now the sample was stabilized at −4.3°C, so that even fewer crystals were observed in the field of view. After stabilization, the samples were supercooled with 0.2°C per minute which forced the crystals to shape in presence of the TIP proteins or type-I AFP purified from winter flounder (wfAFP)^36^ or PBS+ buffer only. Samples were monitored with 1 second intervals using a Nikon 50x ELWD objective. Stills that were used in the figures were taken after 1 minute of supercooling at −4.5°C (full views in **Fig. S8**).

### Crystallization

Crystallization samples were prepared by concentrating TIP-99_a_ protein to 20 mg/mL in PBS+. All crystallization experiments were conducted using the sitting drop vapor diffusion method. Crystallization trials were set up in 200 nL drops using the 96-well plate format at 20 °C. Crystallization plates were set up using a Mosquito from SPT Labtech, then imaged using UVEX microscopes and UVEX PS-600 from JAN Scientific. Diffraction quality crystals formed in 0.02M 1,6-hexanediol, 0.02M 1-butanol, 0.02M 1,2-propanediol, 0.02M 2-propanol, 0.02M 2-propanol, 0.02M 1,4-butanediol, 0.02M 1,3-propanediol, 0.0466M pH 8.5 Tris (base), 0.0534M pH 8.5 Bicine, 20% v/v PEG 500 MME, and 10% w/v PEG 20,000.

X-ray intensities and data reduction were evaluated and integrated using XDS^37^ and merged/scaled using Pointless/Aimless in the CCP4 program suite^38^. Structure determination and refinement starting phases were obtained by molecular replacement using Phaser using the designed model for the structures. Following molecular replacement, the models were improved using phenix.autobuild^40^; efforts were made to reduce model bias by setting rebuild-in-place to false, and using simulated annealing and prime-and-switch phasing. Structures were refined in Phenix^39^. Model building was performed using COOT^41^. The final model was evaluated using MolProbity^42^. Data collection and refinement statistics are recorded in Table S3. Data deposition, atomic coordinates, and structure factors reported in this paper have been deposited in the Protein Data Bank (PDB), http://www.rcsb.org/ with accession code 8EK4.

## Acknowledgments

This work is supported financially by the VLAG graduate school research fellowship to R.J.dH., the Dutch Research Council to R.P.T. (NWO-VENI 202.220), the European Research Council to I.K.V. (ERC-2020-CoG 101001965) and the Audacious Project at the Institute for Protein Design to N.P.K. The authors thank Chuanbao Zheng for assistance in protein purification. Crystallographic work was conducted at the Advanced Photon Source (APS) Northeastern Collaborative Access Team beamlines, which are funded by the National Institute of General Medical Sciences from the National Institutes of Health (P30 GM124165). This research used resources of the Advanced Photon Source, a U.S. Department of Energy (DOE) Office of Science User Facility operated for the DOE Office of Science by Argonne National Laboratory under Contract No. DE-AC02-06CH11357.

