## Supplementary Information for "De novo designed ice-binding proteins from twist-constrained helices"

**This PDF file includes:**

Supporting text

Figures S1 to S8

Tables S1 to S3

SI References

**Other supporting materials for this manuscript include the following:**

Scripts S1 to S6

Supplementary Information Text

**Computational design details**. To generate helical bundle backbones we adapted a python script from Huang et al. (1) and uncouple major (ω_0_) and minor (ω_1_) twisting of the helices. Such that only straight helices of ω_0_=0 would be generated, with a fixed minor twist of ω_1_ = 100, 99.2, 98.2 or 97.2° per residue. After a hyperparameter scan to find optimal parameters for helix offset, phase, bundle radius we typically combinatorically generated 15,000-25,000 three-helix bundles backbones (see **supplementary Script 1** (params.dat) for bundle parameters). Each bundle was designed using two passes Rosetta fixed backbone layer design sequence (see **supplementary for details** **and scripts** for details) and filtered using the packstat filter (2) with a cutoff of 0.3 to remove designs with poor hydrophobic packing in the protein interior. The designs that passed the filter are scored using the Rosetta REF2015 energy function (3). The top 250-500 scoring designs were selected for loop modelling. For each model Rosetta Remodel (4) using a custom blueprint was executed 500 times to design 2 or 3 residues to connect the 3 helices using the KIC protocol into a single chain bundle (see **supplementary** **Table 2 and scripts** for details). Approximately 30% loops resulted in closure. After loop modeling, designs were scored using REF2015 energy functional and the top 75-150 low scoring designs were selected. Rosetta FastRelax (5) was applied to backbones, and the Cα-RMSD to the unrelaxed model were calculated using PyRosetta (see **supplementary** **scripts** for details). Designs were again scored after FastRelax with REF2015 energy function, and selected by RMSD (<1.5Å) and Rosetta score. Rosetta *ab initio* structure prediction was performed starting from primary sequence (see **supplementary** **scripts** for details). Designs that showed a clear folding funnel towards CA-RMSD < 2.5Å with a deep minima were selected for experimental characterization. All design and energy landscape calculations were performed on the WUR Anunna High Performance Computing cluster.

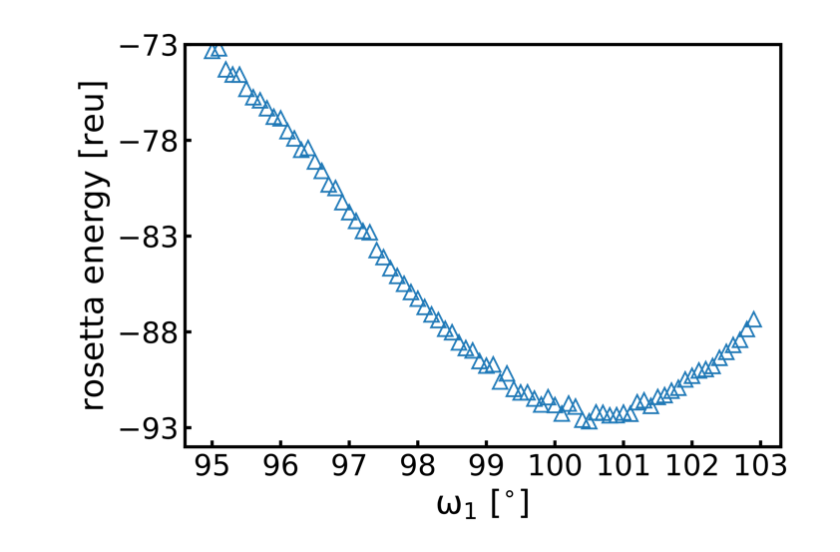

**Fig. S1. Helix stability over minor helical twist.** Single straight (ω_0_=0) alpha-helices of 44 residues were generated using a parametric design script modified from Huang et al. and designed using Rosetta fixed backbone application with amino acid sequence (TAAAAAAAAAA)_4_. Sequences were scored with the Rosetta energy function REF2015 with Rosetta energy units (reu) on the y-axis as a proxy for protein stability: lower reu score means higher stability. A clear minimum is observed at ~100-101° per residue for minor helical twisting of a straight helix.

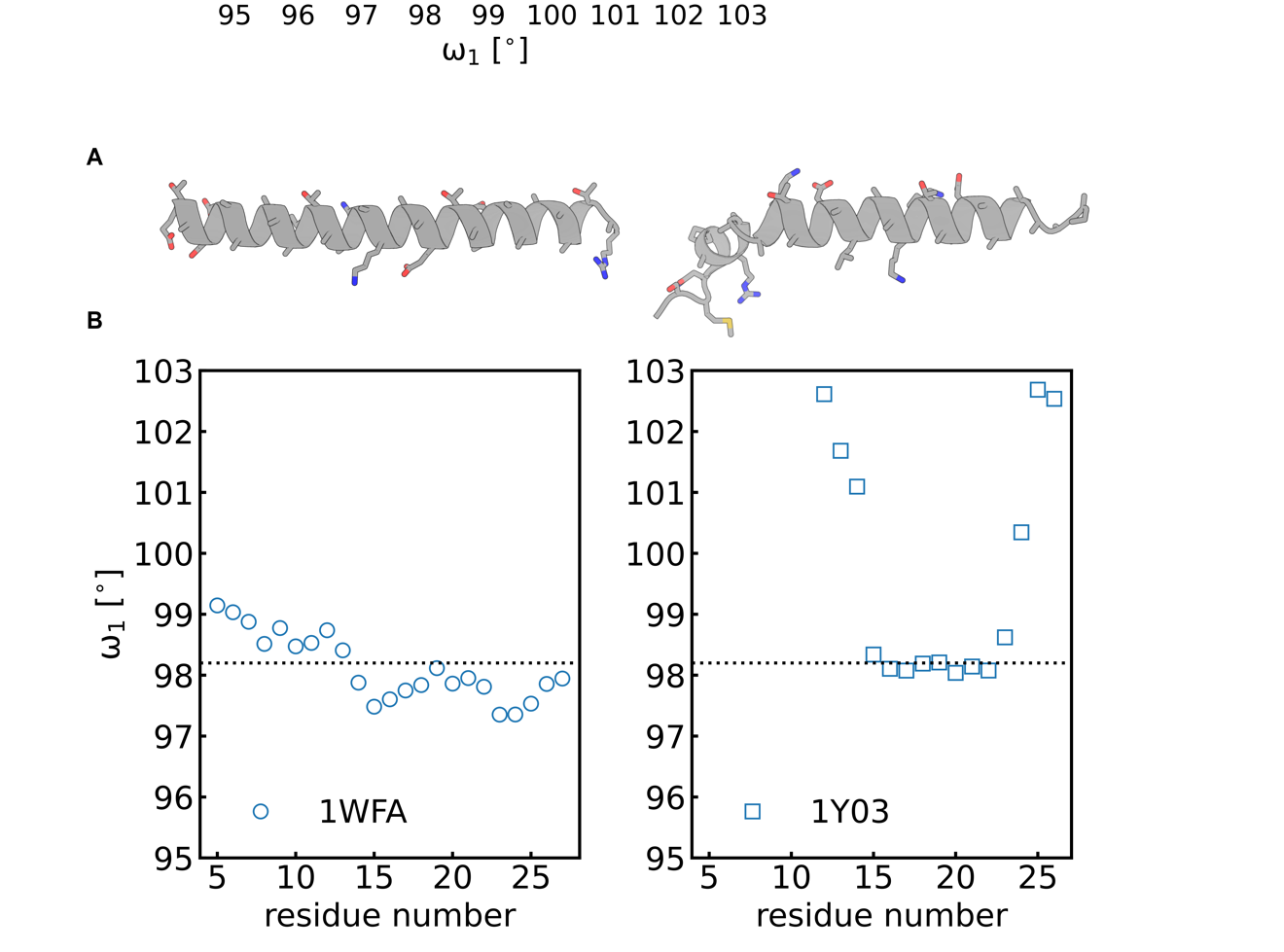

**Fig. S2. Helical twisting for different various type-I ice-binding proteins. A)** experimental structures of type-I ice-binding proteins. *Left:* x-ray crystal structure of wfAFP (pdb accession code: 1WFA) and *right* NMR solution structure of *ss*AFP (pdb accession code: 1Y03, state 1 shown). **B)** mean helical minor twist (ω_1_) over 11 residues calculated with by HELANAL (6) in MDAnalysis (version: 0.20.1) vs. residue number. Dashed line shows expected ice-binding required twisting of ω_1_ = 98.2° per residue which leads to alignment of threonines. For *ss*AFP the N and C termini deviate from 98.2° per residue helical twist, likely due to flexibility.

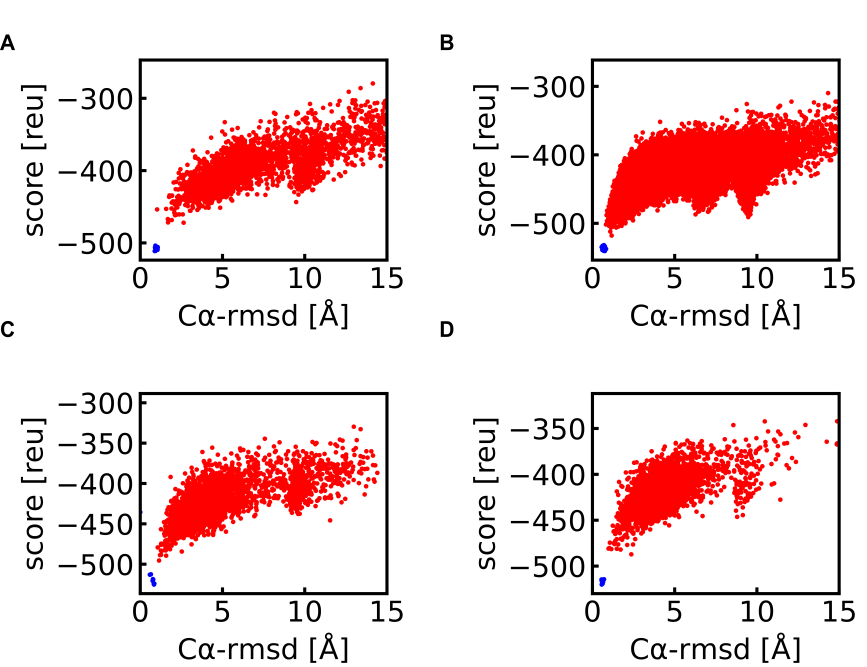

**Fig. S3. Energy landscape of TIP designs. A)** TIP-97, **B)** TIP-98, **C)** TIP-99_a_, **D)** TIP-99_b_, . Score on y-axis shows the rosetta energy of the model in Rosetta energy units [reu] using the energy function REF2015(3) after folding from the extended chain using Rosetta AbinitioRelax simulations, and x-axis shows the Cα-RMSD in [Å] relative to the design model. Each red point represents an individual folding simulation starting with Rosetta AbinitioRelax. Blue points reresent a Rosetta Relaxed model starting from the design model. Funnel-like plots which converge to a single deep minimum near Cα-RMSD< 2.5Å likely demonstrate accurate folding of the design model.

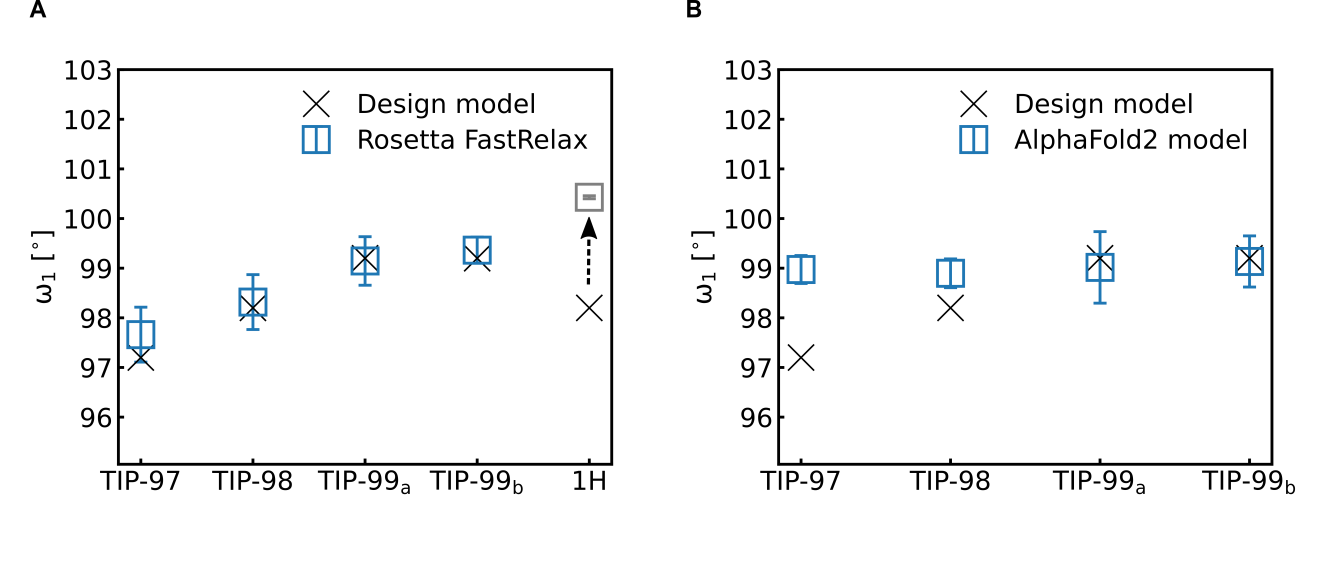
Fig. S4. Comparison helical twisting of ice-binding helix after flexible backbone Rosetta Relax. Y-axis shows the average helical minor twist (ω_1_) of the ice-binding helix calculated by running average over 11 residues using HELANAL (6) in MDAnalysis (version: 0.20.1) A) shows ω_1_ for all characterized designs TIP-99_a_, TIP-99_b_, TIP-98 and TIP-97 after Rosetta Relax with flexible backbone. For all designs the twist of the ice-binding helix is maintained after relaxing with flexible backbone except for 1H which is a single alpha-helix ω_1_=98.2° per residue with sequence (TAAAAAAAAAA)_4_. As expected the 1H helix under-twisting rewinds to the optimal helix twisting of ω_1_=100° per residue because it is not constrained by supporting helices. B) shows ω_1_ of the highest confidence (highest pLDTT ranking) AlphaFold2 model prediction. TIP-97 and TIP-98 show some deviation from the target twist the AF2 predictions. Error bars show standard deviation and X shows the designed ω_1_.

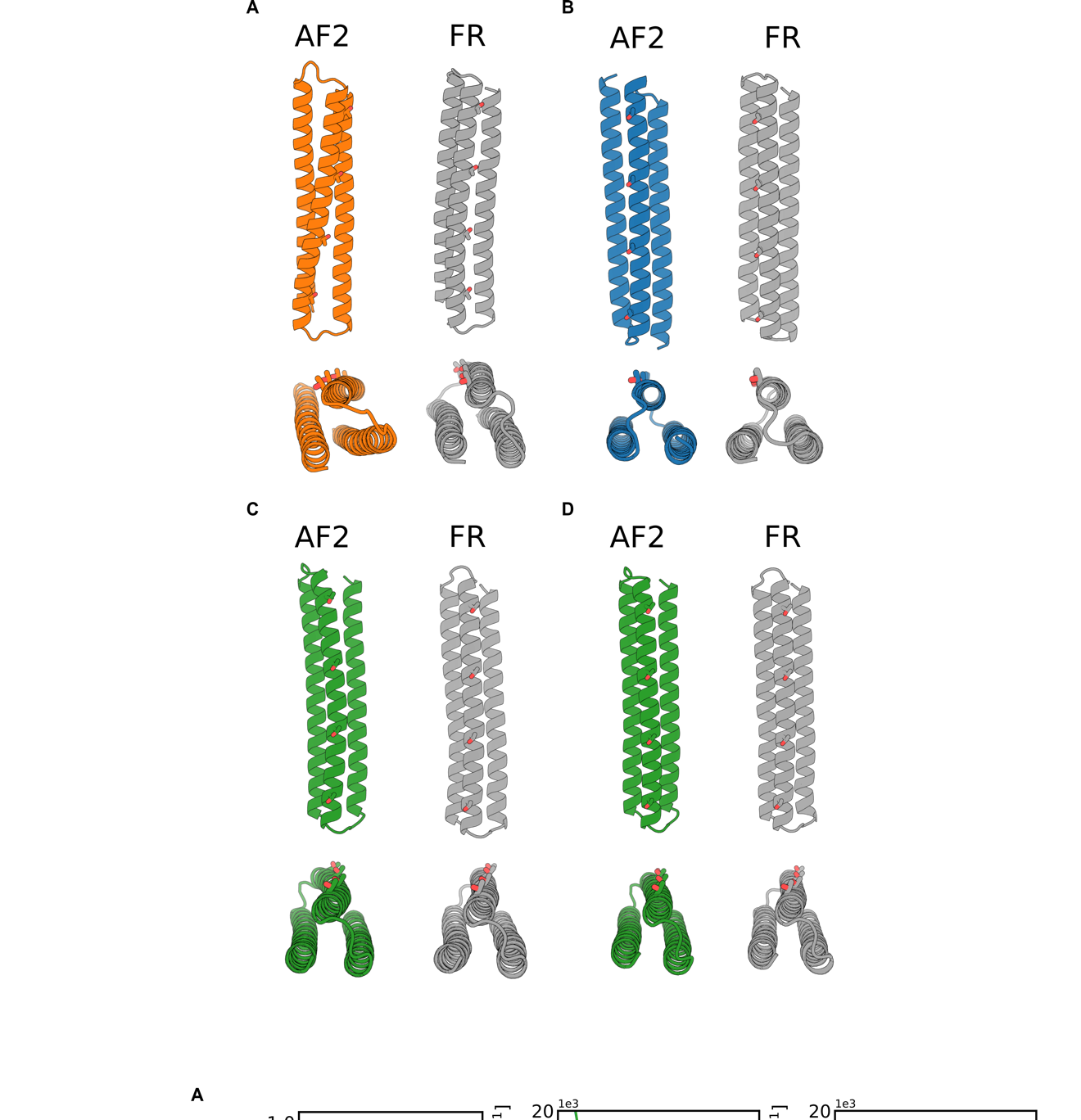

**Fig. S5.** **AlphaFold2 predictions compared to Rosetta FastRelax models.** Cartoons of **A)** TIP-97, **B)** TIP-98, **C)** TIP-99_a_, **D)** TIP-99_b_. AlphaFold2 (AF2) highest ranking pLDTT models (colored) predicted by AlphaFold 2 (7–11) are shown and compared to Rosetta FastRelax (FR) models (grey). Threonine residues on the ice-binding helix are shown as sticks. Noteably: TIP-97 is predicted by AF2 as a bundle with nonzero ω_0_ major helix twist. And TIP-98 is predicted by AF2 to maintain the ω_1_=98.2° of the ice-binding helix, but has a different topology where the two stabilizing helices are reversed in order. TIP-99_a_ and TIP-99_b_ match the Rosetta relaxed structures well both in topology and twisting of the ice-binding helix.

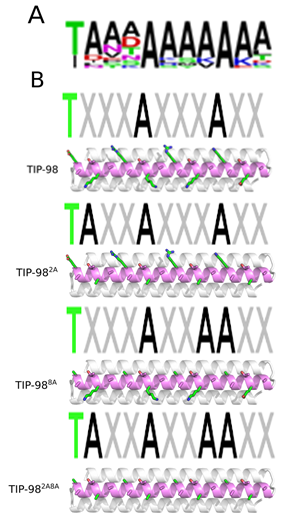

**Fig. S6.** Logo diagram for 11-mer consensus motif computed from sequences of the type-I helical ice-binding proteins *wf*AFP and *ss*AFP. The dominant motif is TAAAAAAAAAA.

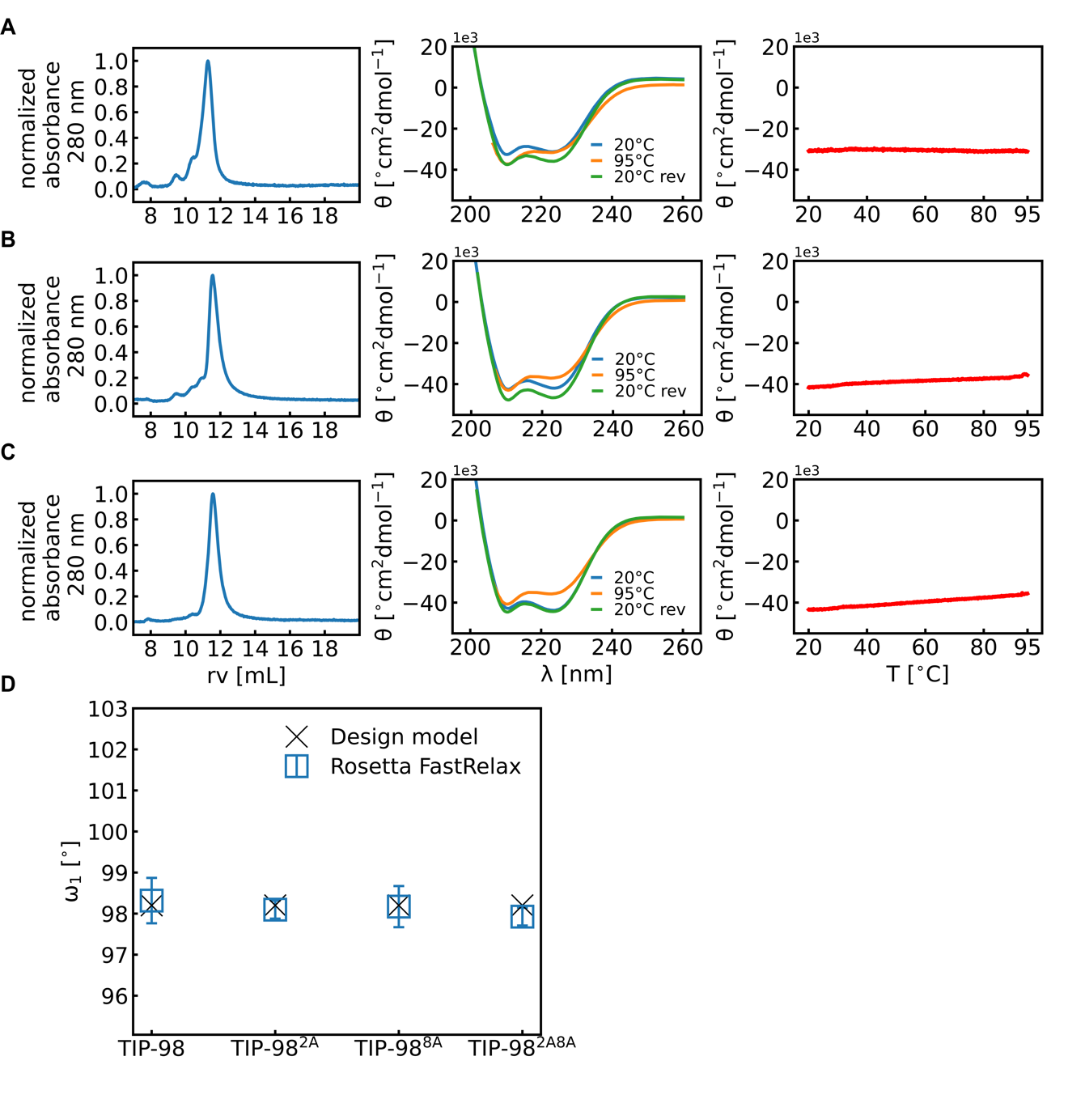

**Fig. S7. Experimental characterization TIP-98 variants with expanded ice-binding motif.** A) TIP-98^2A^ B) TIP-98^8A^ C) TIP-98^2A8A^ Left: size-exclusion chromatography on a Superdex 75 10/300 column in PBS+ (10 mM phosphate pH 7.4 + 300 mM NaCl), demonstrating proteins elute at the expected retention volume (rv) of ~12 mL, corresponding to a 17 kDa globular protein. *Middle*: circular dichroism spectra. Molar ellipticity (θ) versus wavelength (λ) at 20°C, 95°C and 20°C reversed (20°C rev) demonstrating alpha-helical secondary structure and high thermal melt with T_m_ > 95°C, except for TIP-97. Right: molar ellipticity (θ) at 222 nm during temperature ramp from 20°C to 95°C (1°C/min). D) Average helical minor twist (ω_1_) of the ice-binding helix calculated by running average over 11 residues using HELANAL (6) after Rosetta Relax with flexible backbone.

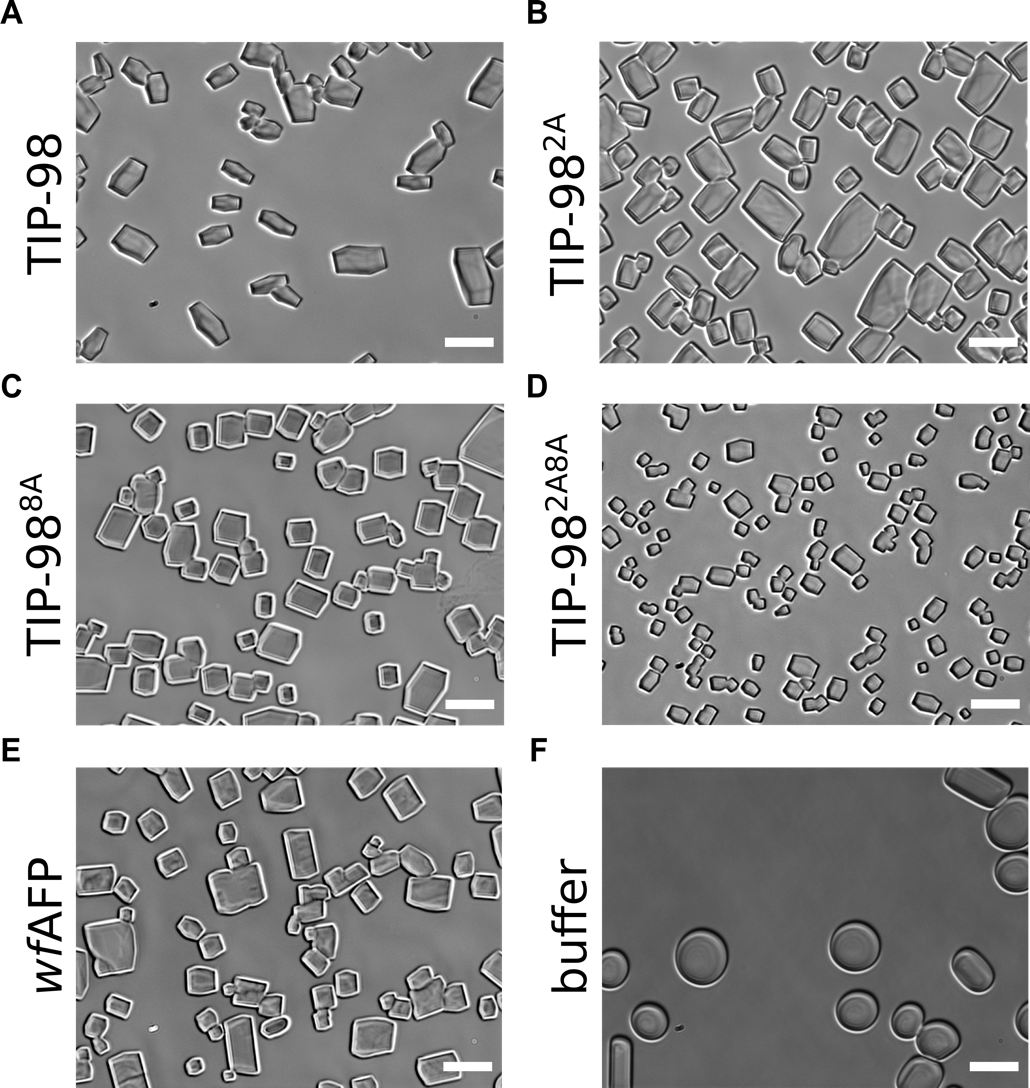

**Fig. S8. Full views of ice shaping experiment.** Images of blunt-end ice-crystals after 1 minute of supercooling at -4.5˚C with 0.2˚C per minute in buffer PBS+ (10 mM phosphate pH 7.4 + 300 mM NaCl) at 50 μM protein concentration. Scale bar: 50 μm.

**Table. S1. proteins sequences of experimentally characterized structures.**

Capitalized sequence is the designed sequence. N-terminally a methionine is added and C-terminally a short serine-glycine linker followed by a tryptophan for extinction at 280nm and a histidine tag for affinity purification. For the variants of TIP-98^2A^, TIP-98^8A^, TIP-98^2A8A^, the alanine mutations are underlined.

| Design name | Protein sequence |
| --- | --- |
| TIP-99_a_ | mEEEAKKKIDDLLTKARREVKKAIKTAREVAKRASKKIEELERRNEDKEAAATKMEAILRAVKTTMKALIEALRTQMKAAAKAMKTIVKAEPESEELKKKVEDAIKDMRRLVEEAIREMEKLARELEKQAREAQKRTsggwhhhhhh |
| TIP-99_b_ | mEEEVREKLKRMEKKFDDSLEKAERKIREIIKEAEKKLKTLKKRNGPYEAVVTTLRAILKAVETKIRAIIKALKTELDALIKAMETILKAHDKNDELKKEVEDIIKKMRDKLTKLIRKAKELLDRLKKKAKKVQDETsggwhhhhhh |
| TIP-98 | mEEEALKKLKDTVKEALKRLKELVDRALKKLKETVKRAEEKLKKLGKDEATITKAKAKLRAIVTKAEADLRALVTKAEAKLKAIVTEASANGVSEEALERLERILREALKRLKKILKEALERLKKILKTAEERLDRNsggwhhhhhh |
| TIP-97 | mKEEFEKKARTLYKRFKTEMDRRITTAERQLTKTARKILTDLRRKKDGGEADLTKAKATLKAQITTIIARFRARATKLIAQINAELTKDVAKMPVDEEMQKRLEKILKTLEETARRLLKTMTTRMQQTLEKIITTMDREsggwhhhhhh |
| TIP-98^2A^ | mEEEALKKLKDTVKEALKRLKELVDRALKKLKETVKRAEEKLKKLGKDEATITAAKAKLRAIVTAAEADLRALVTAAEAKLKAIVTAASANGVSEEALERLERILREALKRLKKILKEALERLKKILKTAEERLDRNssgwhhhhhh |
| TIP-98^8A^ | mEEEALKKLKDTVKEALKRLKELVDRALKKLKETVKRAEEKLKKLGKDAATITKAKAKLAAIVTKAEADLAALVTKAEAKLAAIVTEASANGVSEEALERLERILREALKRLKKILKEALERLKKILKTAEERLDRNssgwhhhhhh |
| TIP-98^2A8A^ | mEEEALKKLKDTVKEALKRLKELVDRALKKLKETVKRAEEKLKKLGKDAATITAAKAKLAAIVTAAEADLAALVTAAEAKLAAIVTAASANGVSEEALERLERILREALKRLKKILKEALERLKKILKTAEERLDRNssgwhhhhhh |

**Table. S2. AlphaFold 2 per residue confidence scores (pLDDT) for the 5 parameter models.** A higher pLDDT score is better. For each TIP design, the highest ranking pLDTT model is underlined.

| **Design** | **Model 1** | **Model 2** | **Model 3** | **Model 4** | **Model 5** |
| --- | --- | --- | --- | --- | --- |
| **TIP-97** | 83.32 | 84.60 | 81.87 | 80.54 | 79.10 |
| **TIP-98** | 86.07 | 91.01 | 92.79 | 89.87 | 89.49 |
| **TIP-99_a_** | 90.46 | 92.76 | 91.29 | 90.47 | 89.63 |
| **TIP-99_b_** | 90.88 | 92.14 | 92.20 | 91.13 | 90.30 |

**Table S3. Crystallographic data collection and refinement statistics**

| Design | TIP-99_a_ |
| --- | --- |
| Resolution range | 49.94 - 2.35 (2.434 - 2.35) |
| Space group | P 21 21 21 |
| Unit cell | 54.582 67.427 74.326 90 90 90 |
| Total reflections | 157388 (16229) |
| Unique reflections | 11928 (1172) |
| Multiplicity | 13.2 (13.8) |
| Completeness (%) | 99.62 (100.00) |
| Mean I/sigma(I) | 13.80 (1.55) |
| Wilson B-factor | 60,27 |
| R-merge | 0.1145 (1.379) |
| R-meas | 0.1193 (1.431) |
| R-pim | 0.03301 (0.3815) |
| CC1/2 | 0.999 (0.753) |
| CC* | 1 (0.927) |
| Reflections used in refinement | 11887 (1173) |
| Reflections used for R-free | 553 (49) |
| R-work | 0.2610 (0.3181) |
| R-free | 0.3189 (0.3946) |
| CC (work) | 0.964 (0.575) |
| CC (free) | 0.930 (0.316) |
| macromolecules | 2165 |
| RMS(bonds) | 0,002 |
| RMS(angles) | 0,39 |
| Ramachandran favored (%) | 97,74 |
| Ramachandran allowed (%) | 2,26 |
| Ramachandran outliers (%) | 0 |
| Rotamer outliers (%) | 0,43 |
| Average B-factor | 83,35 |
| macromolecules | 83,35 |

***Script S1. Parametric bundle backbone scripts***

Example of parameters (params.dat) used to generate ~17,500 TIP-99 helices. params.dat adapted from Huang et al. (1) Python (generate_helix.py) script adapted from Huang et al.(1) to generate helices edited such that in the generation of helices major superhelical (ω_0_) and minor helix twist (ω_1_) are de-coupled, and ω_1_=98.2° per residue for all helices.

```
params.dat
TIP-99         # base name of output structures
44             # number of residues per helix
6.0 7.0 4      # supercoil radius range (low, high, number of samples)
3              # number of helices
3              # number of helices in asymmetric unit
1 -1 1         # orientation of each helix in asymmetric unit
200 250 9      # phase of first helix (low, high, number of samples)
170 200 3      # phase of second helix (low, high, number of samples)
200 250 9      # phase of third helix (low, high, number of samples)
-3.0 3.0 6     # z-offset of first helix (low, high, number of samples)
0.0 0.0 1      # z-offset of second helix (low, high, number of samples)
-3.0 3.0 3     # z-offset of third helix (low, high, number of samples)
A B C          # chain names
0 1 2          # chain order
```

generate_helix.py (for generation of helices with ω_1_=99.2° per residue)

```
###################Start of script##############################
#!/usr/bin/env python
from os import system,popen
import string
from sys import argv
import math
from math import sin,cos,sqrt
def  isint   (x): return type(x) is int
def  isfloat (x): return type(x) is float
def isnum  (x): return isint(x) or isfloat(x)
def  isiter  (x): return hasattr(x,"__iter__")
def  islist  (x): return type(x) is list
def istuple(x): return type(x) is tuple
def isvec  (x): return type(x) is Vec
def ismat  (x): return type(x) is Mat
def isxform(x): return type(x) is Xform
class Vec(object):
        """a Vector like xyzVector<Real> in rosetta
        >>> v = Vec(1,2,3)
        >>> print v, 10*v
        (1.000000,2.000000,3.000000) (10.000000,20.000000,30.000000)
        multiplication is a dot prod at the moment
        >>> v*v
        14.0
        >>> assert Vec(1,0,-0) == Vec(1,-0,0)
        """
        def __init__(self,x=0.0,y=None,z=None):
                if y is None:
                        if isnum(x):
                                self.x,self.y,self.z = (float(x),)*3
                        elif isvec(x):
                                self.x,self.y,self.z = x.x,x.y,x.z
                        elif isiter(x):
                                i = iter(x)
                                self.x,self.y,self.z = i.next(),i.next(),i .next()
                        else: raise NotImplementedError
                elif z is not None:
                        assert isnum(x) and isnum(y) and isnum(z)
                        self.x,self.y, self .z = float(x),float (y), float (z)
                else: raise NotImplementedError
                assert isfloat(self .x)
                assert isfloat(self .y)
                assert isfloat(self .z)
        def dot(u,v):     assert isvec(v); return u.x*v.x+u.y*v.y+u.z*v.z
        def normdot(u,v): assert isvec(v); return min(1.0,max(-1.0,u.dot(v)/u.length()/v.length()))
        def angle(u,v):
                assert isvec(v)
                d = u.normdot(v)
                if d > 1.0-EPS: return 0.0;
                if d < EPS-1.0: return pi
                return acos(d)
        def angle_degrees(u,v): return degrees(u.angle(v))
        def lineangle(u,v):
                assert isvec(v);
                if u.length() < SQRTEPS or v.length < SQRTEPS: return 0.0
                ang = abs(acos( u.normdot(v) ))
                return ang if ang < pi/2.0 else pi-ang
        def lineangle_degrees(u,v):
                if isvec(v): return degrees(u.lineangle(v))
                raise NotImplementedError
        def length(u):         return sqrt(u.dot(u))
        def length_squared(u): return      u.dot(u)
        def distance(u,v):         assert isvec(v); return (u-v).length()
        def distance_squared(u,v): assert isvec(v); return (u-v).length_squared()
        def cross(u,v): assert isvec(v); return Vec(u.y*v.z-u.z*v.y,u.z*v.x-u.x*v.z,u.x*v.y-u.y*v.x)
        def __mul__(u,a):
                if   isnum(a): return Vec(u.x*a,u.y*a,u.z*a)
                elif isvec(a): return u.dot(a)
                else: return a.__rmul__(u)
        def __rmul__(u,a): return u*a
        def __add__(u,v):
                if isvec(v): return Vec(u.x+v.x,u.y+v.y,u.z+v.z)
                return v.__radd__(u)
        def __sub__(u,v):
                if isvec(v): return Vec(u.x-v.x,u.y-v.y,u.z-v.z)
                return v.__rsub__(u)
        def __neg__(u): return Vec(-u.x,-u.y,-u.z)
        def __div__(u,a): return u*(1.0/a)
        def __str__(self): return "(%f,%f,%f)"%(self.x,self.y,self.z)
        def __repr__(self): return "Vec( %f, %f, %f )"%(self.x,self.y,self.z)
        def normalize(u):
                l = u.length()
                u.x /= l
                u.y /= l
                u.z /= l
        def normalized(u):
                v = Vec(u)
                v.normalize()
                return v
        def outer(u,v):
                assert isvec(v)
                return Mat( u.x*v.x, u.x*v.y, u.x*v.z,
                                                u.y*v.x, u.y*v.y, u.y*v.z,
                                                u.z*v.x, u.z*v.y, u.z*v.z      )
        def __eq__(self,other):
                assert isvec(other)
                return ( abs(self.x-other.x) < EPS and
                                        abs(self.y-other.y) < EPS and
                                        abs(self.z-other.z) < EPS )
        def rounded(self,sd):
                return Vec(round(self.x,sd), round(self.y,sd), round(self.z,sd) )
        def unit(v):
                if   abs(v.x) > SQRTEPS: return v/v.x
                elif abs(v.y) > SQRTEPS: return v/v.y
                elif abs(v.z) > SQRTEPS: return v/v.z
        def __len__(v):
                return 3
        def abs(v):
                return Vec(abs(v.x),abs(v.y),abs(v.z))
        def __getitem__(v,i):
                if i is 0: return v.x
                if i is 1: return v.x
                if i is 2: return v.x
                raise IndexError
        def tuple(v):
                return (v.x,v.y,v.z)
        def key(v):
                return v.abs().unit().rounded(6).tuple()
Ux = Vec(1,0,0)
Uy = Vec(0,1,0)
Uz = Vec(0,0,1)
V0 = Vec(0,0,0)
def randvec(n=None):
        if n is None: return Vec(gauss(0,1),gauss(0,1),gauss(0,1))
        return [Vec(gauss(0,1),gauss(0,1),gauss(0,1)) for i in range(n)]
def randveccube(r=1.0):
        return Vec(uniform(-1,1),uniform(-1,1),uniform(-1,1))*r
def randvecball(r=1.0):
        v =randveccube(r)
        while v.length_squared() > r*r: v = randveccube(r)
        return v
def randnorm(n=None):
        """
        >>> assert abs(randnorm().length()-1.0) < 0.0000001
        """
        if n is None: return randvec().normalized()
        return (randvec().normalized() for i in range(n))
def coplanar(x1,x2,x3,x4):
        """
        >>> u,v,w = randvec(3)
        >>> a,b,c = (gauss(0,10) for i in range(3))
        >>> assert     coplanar(u, v, w, u + a*(u-v) + b*(v-w) + c*(w-u) )
        >>> assert not coplanar(u, v, w, u + a*(u-v) + b*(v-w) + c*(w-u) + randvec().cross(u-v) )
        """
        return abs((x3-x1).dot((x2-x1).cross(x4-x3))) < SQRTEPS
def rmsd(l,m):
        """
        >>> l,m = randvec(6),randvec(6)
        >>> rmsd(l,l)
        0.0
        """
        rmsd = 0.0
        for u,v in izip(l,m):
                rmsd += u.distance_squared(v)
        return sqrt(rmsd)
class Mat(object):
        """docstring for Mat
        >>> m = Mat(2,0,0,0,1,0,0,0,1)
        >>> print m
        Mat[ (2.000000,0.000000,0.000000), (0.000000,1.000000,0.000000), (0.000000,0.000000,1.000000) ]
        >>> print m*m
        Mat[ (4.000000,0.000000,0.000000), (0.000000,1.000000,0.000000), (0.000000,0.000000,1.000000) ]
        >>> print Mat(*range(1,10)) * Mat(*range(10,19))
        Mat[ (84.000000,90.000000,96.000000), (201.000000,216.000000,231.000000), (318.000000,342.000000,366.000000) ]
        >>> assert Mat(0.0,1.0,2.0,3,4,5,6,7,8) == Mat(-0,1,2,3,4,5.0,6.0,7.0,8.0)
        >>> print Mat(100,2,3,4,5,6,7,8,9).det()
        -297.0
        >>> m = Mat(100,2,3,4,5,6,7,8,9)
        >>> assert m * ~m == Imat
        """
        def __init__(self, xx=None, xy=None, xz=None, yx=None, yy=None, yz=None, zx=None, zy=None, zz=None):
                super(Mat, self).__init__()
                if xx is None: # identity default
                        self.xx, self.xy, self.xz = 1.0,0.0,0.0
                        self.yx, self.yy, self.yz = 0.0,1.0,0.0
                        self.zx, self.zy, self .zz = 0.0,0.0,1.0
                elif xy is None and ismat(xx):
                        self.xx, self.xy, self.xz = xx.xx, xx.xy, xx.xz
                        self.yx, self.yy, self.yz = xx.yx, xx.yy, xx.yz
                        self.zx, self.zy, self .zz = xx.zx, xx.zy, xx.zz
                elif yx is None and isvec(xx) and isvec(xy) and isvec(xz):
                        self.xx, self.xy, self.xz = xx.x, xy.x, xz.x
                        self.yx, self.yy, self.yz = xx.y, xy.y, xz.y
                        self.zx, self.zy, self .zz = xx.z, xy.z, xz.z
                elif isnum(xx):
                        self.xx, self.xy, self.xz = float(xx), float(xy), float(xz)
                        self.yx, self.yy, self.yz = float(yx), float(yy), float(yz)
                        self.zx, self.zy, self .zz = float(zx), float(zy), float(zz)
                else:
                        raise NotImplementedError
                assert isfloat(self .xx) and isfloat(self .xy) and isfloat(self .xz)
                assert isfloat(self .yx) and isfloat(self .yy) and isfloat(self .yz)
                assert isfloat(self .zx) and isfloat( self .zy) and isfloat( self .zz)
        def row(m,i):
                assert isint(i)
                if   i is 0: return Vec(m.xx,m.xy,m.xz)
                elif i is 1: return Vec(m.yx,m.yy,m.yz)
                elif i is 2: return Vec(m.zx,m.zy,m.zz)
                else: assert 0 <= i and i <= 2
        def col(m,i):
                assert isint(i)
                if   i is 0: return Vec(m.xx,m.yx,m.zx)
                elif i is 1: return Vec(m.xy,m.yy,m.zy)
                elif i is 2: return Vec(m.xz,m.yz,m.zz)
                else: assert 0 <= i and i <= 2
        def rowx(m): return m.row(0)
        def rowy(m): return m.row(1)
        def rowz(m): return m.row(2)
        def colx(m): return m.col(0)
        def coly(m): return m.col(1)
        def colz(m): return m.col(2)
        def __invert__(m):
                """
                >>> from random import random
                >>> for i in range(10):
                ...     m = random()*Mat(random(),random(),random(),random(),random(),random(),random(),random(),random())
                ...     assert ~m*m == Imat
                ...     assert m*~m == Imat
                """
                return Mat(m.zz*m.yy-m.zy*m.yz, -(m.zz*m.xy-m.zy*m.xz),m.yz*m.xy-m.yy*m.xz,
                         -(m.zz*m.yx-m.zx*m.yz),m.zz*m.xx-m.zx*m.xz,-(m.yz*m.xx-m.yx*m.xz),
                          m.zy*m.yx-m.zx*m.yy,-(m.zy*m.xx-m.zx*m.xy),m.yy*m.xx-m.yx*m.xy)/ m.det()
        def __mul__(m,rhs):
                if   isnum(rhs): return Mat( rhs*m.xx, rhs*m.xy, rhs*m.xz, rhs*m.yx, rhs*m.yy, rhs*m.yz, rhs*m.zx, rhs*m.zy, rhs*m.zz )
                elif isvec(rhs): return Vec( m.rowx()*rhs, m.rowy()*rhs, m.rowz()*rhs )
                elif ismat(rhs): return Mat( m.rowx()*rhs.colx(), m.rowx()*rhs.coly(), m.rowx()*rhs.colz(),
                                             m.rowy()*rhs.colx(), m.rowy()*rhs.coly(), m.rowy()*rhs.colz(),
                                             m.rowz()*rhs.colx(), m.rowz()*rhs.coly(), m.rowz()*rhs.colz() )
                else: return rhs.__rmul__(m)
        def __rmul__(m,v):
                if   isnum(v): return m*v
                elif isvec(v): return Vec( m.colx()*v, m.coly()*v, m.colz()*v )
        def __div__(m,v): return m*(1/v)
        def __add__(m,v):
                if   isnum(v): return Mat(v+m.xx,v+m.xy,v+m.xz,v+m.yx,v+m.yy,v+m.yz,v+m.zx,v+m.zy,v+m.zz)
                elif ismat(v): return Mat(v.xx+m.xx,v.xy+m.xy,v.xz+m.xz,v.yx+m.yx,v.yy+m.yy,v.yz+m.yz,v.zx+m.zx,v.zy+m.zy,v.zz+m.zz)
                else: return v.__radd__(m)
        def __sub__(m,v): return m + -v
        def __neg__(m): return m * -1
        def __str__(m): return "Mat[ %s, %s, %s ]" % (str(m.rowx()),str(m.rowy()),str(m.rowz()))
        def __repr__(m): return "Mat( %s, %s, %s )" % (repr(m.colx()),repr(m.coly()),repr(m.colz()))
        def transpose(m):
                m = Mat( m.xx, m.yx, m.zx, m.xy, m.yy, m.zy, m.xz, m.yz, m.zz )
        def transposed(m): return Mat( m.xx, m.yx, m.zx, m.xy, m.yy, m.zy, m.xz, m.yz, m.zz )
        def det(m):
                 #   a11  (a33  a22- a32  a23)- a21 ( a33  a12- a32  a13)+ a31(  a23  a12- a22  a13)
                return m.xx*(m.zz*m.yy-m.zy*m.yz)-m.yx*(m.zz*m.xy-m.zy*m.xz)+m.zx*(m.yz*m.xy-m.yy*m.xz)
        def trace(m): return m.xx+m.yy+m.zz
        def add_diagonal(m,v): return Mat( v.x+m.xx, m.xy, m.xz, m.yx, v.y+m.yy, m.yz, m.zx, m.zy, v.z+m.zz )
        def is_rotation(m): return (m.colx().isnormal() and m.coly().isnormal() and m.colz().isnormal() and
                                  m.rowx().isnormal() and m.rowy().isnormal() and m.rowz().isnormal()   )
        def __eq__(self,other): return ( abs(self.xx-other.xx) < EPS and
                                        abs(self.xy-other.xy) < EPS and
                                        abs(self.xz-other.xz) < EPS and
                                        abs(self.yx-other.yx) < EPS and
                                        abs(self.yy-other.yy) < EPS and
                                        abs(self.yz-other.yz) < EPS and
                                        abs(self.zx-other.zx) < EPS and
                                        abs(self.zy-other.zy) < EPS and
                                        abs(self.zz-other.zz) < EPS )
        def rotation_axis(R):
                """
                >>> axis ,ang  = randnorm(),uniform(-pi,pi)
                >>> axis2,ang2 = rotation_matrix(axis,ang).rotation_axis()
                >>> assert abs( abs(ang) - abs(ang2) ) < EPS
                >>> assert axis == axis2 * copysign(1,ang*ang2)
                """
                cos_theta = sin_cos_range((R.trace()-1.0)/2.0);
                if cos_theta > -1.0+EPS and cos_theta < 1.0-EPS:
                        x = ( 1.0 if R.zy > R.yz else -1.0 ) * sqrt( max(0.0, ( R.xx - cos_theta ) / ( 1.0 - cos_theta ) ) )
                        y = ( 1.0 if R.xz > R.zx else -1.0 ) * sqrt( max(0.0, ( R.yy - cos_theta ) / ( 1.0 - cos_theta ) ) )
                        z = ( 1.0 if R.yx > R.xy else -1.0 ) * sqrt( max(0.0, ( R.zz - cos_theta ) / ( 1.0 - cos_theta ) ) )
                        theta = acos( cos_theta );
                        assert abs( x*x + y*y + z*z - 1 ) <= 0.01
                        return Vec(x,y,z),theta
                elif cos_theta >= 1.0-EPS: return Vec(1.0,0.0,0.0),0.0
                else:
                        nnT = (R+Imat)/2.0
                        x,y,z = 0.0,0.0,0.0;
                        if nnT.xx > EPS:
                                x = sqrt( nnT.xx )
                                y = nnT.yx / x
                                z = nnT.zx / x
                        elif nnT.yy > EPS:
                                x = 0
                                y = sqrt(nnT.yy)
                                z = nnT.zy / y
                        else:
                                assert( nnT.zz > EPS );
                                x = 0
                                y = 0
                                z = sqrt( nnT.zz )
                        assert abs( x*x + y*y + z*z - 1.0 ) <= 0.01
                        return Vec( x, y, z ),pi
        def euler_angles(self):
                FLOAT_PRECISION = 1e-5
                if self.zz >= 1 - FLOAT_PRECISION:
                        e1 = math.acos( sin_cos_range( self.xx ) )
                        e2 = 0.0
                        e3 = 0.0
                        return Vec(e1,e2,e3)
                if self.zz <= -1 + FLOAT_PRECISION:
                        e1 = math.acos( sin_cos_range( self.xx ) )
                        e2 = 0.0
                        e3 = math.pi
                        return Vec(e1,e2,e3)
                pos_sin_theta = math.sqrt( 1 - self.zz*self.zz ) # sin2theta = 1 - cos2theta.
                #  two values are  possible   here : my convention   is  to use  positive   theta   only .
                #  corresponding theta  between   [0, pi /2] -> [0,90]  since   st  > 0
                #  and asin returns value  between  [-pi /2, pi /2]
                e3 = math.asin( pos_sin_theta )
                #  decide whether the  actual   positive   theta   is   between  [ pi /2, pi [  using   the   value  of cos ( theta )
                #  which happens to be the matrix  element   self . zz (and  is   thus   signed ) .
                if self.zz < 0:
                        e3 = math.pi - e3
                # e1 = math.atan2(  -UU(1,3), UU(2,3) )   #  between -Pi and Pi -> [-180,180]
                # e2 = math.atan2( UU(3,1), UU(3,2) )     #  between -Pi and Pi -> [-180, 180]
                #  this is atan( sin_phi * c , cos_phi * c  ) as  opposed  to Alex ' s atan ( -sin_phi * c, -cos_phi * c ) .
                e1 = math.atan2( self.zx, -self.zy )
                e2 = math.atan2( self.xz,  self.yz )
                return Vec(e1,e2,e3)
        def from_euler_angles(self,euler):
                phi   = euler.x()
                psi   = euler.y()
                theta = euler.z()
                cos_phi   = math.cos( phi )
                sin_phi   = math.sin( phi )
                cos_psi   = math.cos( psi )
                sin_psi   = math.sin( psi )
                cos_theta = math.cos( theta )
                sin_theta = math.sin( theta )
                self.xx =  cos_psi * cos_phi - cos_theta * sin_phi * sin_psi
                self.xy =  cos_psi * sin_phi + cos_theta * cos_phi * sin_psi
                self.xz =  sin_psi * sin_theta
                self.yx = -sin_psi * cos_phi - cos_theta * sin_phi * cos_psi
                self.yy = -sin_psi * sin_phi + cos_theta * cos_phi * cos_psi
                self.yz =  cos_psi * sin_theta
                self.zx =  sin_theta * sin_phi
                self.zy = -sin_theta * cos_phi
                self.zz =        cos_theta
                return self
Imat = Mat(1,0,0,0,1,0,0,0,1)
class Xform(object):
        """Coordinate frame like rosetta Xform, behaves also as a rosetta Stub
        >>> x = Xform(R=Imat,t=Uz)
        >>> print x
        Xform( Mat[ (1.000000,0.000000,0.000000), (0.000000,1.000000,0.000000), (0.000000,0.000000,1.000000) ], (0.000000,0.000000,1.000000) )
        >>> assert (x*x) == Xform(R=Imat,t=2*Uz)
        >>> x = Xform(R=rotation_matrix_degrees(Vec(1,0,0),90.0),t=Vec(0,0,0))
        >>> print x
        Xform( Mat[ (1.000000,0.000000,0.000000), (0.000000,0.000000,-1.000000), (0.000000,1.000000,0.000000) ], (0.000000,0.000000,0.000000) )
        >>> assert x*x*x*x == Ixform
        >>> x.t = Ux
        >>> assert x*x*x*x == Xform(R=Imat,t=4*Ux)
        >>> x.t = Uz
        >>> print x
        Xform( Mat[ (1.000000,0.000000,0.000000), (0.000000,0.000000,-1.000000), (0.000000,1.000000,0.000000) ], (0.000000,0.000000,1.000000) )
        >>> assert x               == Xform(R=rotation_matrix_degrees(Ux, 90.0),t=Vec(0, 0,1))
        >>> assert x*x             == Xform(R=rotation_matrix_degrees(Ux,180.0),t=Vec(0,-1,1))
        >>> assert x*x*x           == Xform(R=rotation_matrix_degrees(Ux,270.0),t=Vec(0,-1,0))
        >>> assert x*x*x*x         == Xform(R=rotation_matrix_degrees(Ux,  0.0),t=Vec(0, 0,0))
        >>> assert x*x*x*x*x       == Xform(R=rotation_matrix_degrees(Ux, 90.0),t=Vec(0, 0,1))
        >>> assert x*x*x*x*x*x     == Xform(R=rotation_matrix_degrees(Ux,180.0),t=Vec(0,-1,1))
        >>> assert x*x*x*x*x*x*x   == Xform(R=rotation_matrix_degrees(Ux,270.0),t=Vec(0,-1,0))
        >>> assert x*x*x*x*x*x*x*x == Xform(R=rotation_matrix_degrees(Ux,  0.0),t=Vec(0, 0,0))
        >>> x = Xform(rotation_matrix_degrees(Vec(1,2,3),123),Vec(5,7,9))
        >>> assert ~x *  x == Ixform
        >>> assert  x * ~x == Ixform
        Frames / RTs are interchangable:
        >>> fr = Xform(rotation_matrix_degrees(Vec(1,2,3), 65.64),t=Vec(3,2,1))
        >>> to = Xform(rotation_matrix_degrees(Vec(7,5,3),105.44),t=Vec(10,9,8))
        >>> x = to/fr
        >>> assert to/Ixform ==  to
        >>> assert Ixform/fr == ~fr
        >>> assert (to * ~fr) * fr == to
        >>> assert x * fr == to
        >>> a1 = randnorm()
        >>> b1 = randnorm()
        >>> ang = uniform(0,1)*360.0-180.0
        >>> a2 = rotation_matrix_degrees(a1.cross(randnorm()),ang) * a1
        >>> b2 = rotation_matrix_degrees(b1.cross(randnorm()),ang) * b1
        >>> assert abs(angle(a1,a2) - angle(b1,b2)) < EPS
        >>> xa = Xform().from_two_vecs(a1,a2)
        >>> xb = Xform().from_two_vecs(b1,b2)
        >>> assert xa.tolocal(a1) == xb.tolocal(b1)
        >>> assert xa.tolocal(a2) == xb.tolocal(b2)
        >>> assert ~xa*a1 == ~xb*b1
        >>> assert ~xa*a2 == ~xb*b2
        >>> assert xb/xa*a1 == b1
        >>> assert xb/xa*a2 == b2
        add/sub with Vecs:
        >>> X = randxform()
        >>> u,v = randvec(2)
        >>> assert isxform(u+X) and isxform(X+u) and isxform(u-X) and isxform(X-u)
        >>> assert X*v+u == (u+X)*v
        >>> assert X*(v+u) == (X+u)*v
        >>> assert Xform(u)*X*v == (u+X)*v
        >>> assert X*Xform(u)*v == (X+u)*v
        >>> assert X*v-u == (u-X)*v
        >>> assert X*(v-u) == (X-u)*v
        mul,div with Mats:
        >>> R = randrot()
        >>> assert isxform(R*X) and isxform(X*R)
        >>> assert R*X*u == (R*X)*u == R*(X*u)
        >>> assert X*R*u == (X*R)*u == X*(R*u)
        >>> assert Xform(R)*X*u == Xform(R)*(X*u)
        >>> assert X*Xform(R)*u == X*(Xform(R,V0)*u)
        >>> assert X/X*v == v
        mul/div Xforms:
        >>> Y = randxform()
        >>> assert isxform(X/Y) and isxform(X*Y)
        >>> assert X/Y*v == X*~Y*v
        these don't work yet:
        >>> axis,ang,cen = randnorm(),uniform(-pi,pi),randvec()      #doctest: +SKIP
        >>> X = rotation_around(axis,ang,cen)                        #doctest: +SKIP
        >>> axis2,ang2,cen2 = X.rotation_center()                    #doctest: +SKIP
        >>> assert abs( abs(ang) - abs(ang2) ) < EPS                 #doctest: +SKIP
        >>> assert axis == axis2 * copysign(1,ang*ang2)              #doctest: +SKIP
        >>> print cen                                                #doctest: +SKIP
        >>> print cen2                                               #doctest: +SKIP
        >>> x =                Xform( Mat( Vec( 0.816587, -0.306018, 0.489427 ), Vec( 0.245040, 0.951487, 0.186086 ), Vec( -0.522629, -0.032026, 0.851959 ) ), Vec( 1.689794, 1.535762, -0.964428 ) )
        >>> assert repr(x) == "Xform( Mat( Vec( 0.816587, -0.306018, 0.489427 ), Vec( 0.245040, 0.951487, 0.186086 ), Vec( -0.522629, -0.032026, 0.851959 ) ), Vec( 1.689794, 1.535762, -0.964428 ) )"
        """
        def __init__(self, R=None, t=None):
                super(Xform, self).__init__()
                if isvec(R) and t is None: R,t = Imat,R
                self.R = R if R else Imat
                self.t = t if t else V0
                assert ismat(self.R) and isvec(self.t)
        def from_four_points(s,cen,a,b,c):
                s.t = cen
                e1 = (a-b).normalized()
                e3 = e1.cross(c-b).normalized()
                e2 = e1.cross(e3).normalized()
                s.R = Mat(e1.x,e2.x,e3.x,e1.y,e2.y,e3.y,e1.z,e2.z,e3.z)
                return s
        def from_two_vecs(s,a,b):
                e1 = a.normalized()
                e2 = projperp(a,b).normalized()
                e3 = e1.cross(e2)
                return Xform( Mat(e1.x,e2.x,e3.x,e1.y,e2.y,e3.y,e1.z,e2.z,e3.z),V0)
        def tolocal(s,x): return s.R.transposed() * (x - s.t)
        def toglobal(s,x): return (s.R * x) + s.t
        def __invert__(self):
                R = ~self.R
                t = R * -self.t
                return Xform(R,t)
        def __mul__(X,o):
                if   isvec(o):   return X.R * o + X.t
                elif isxform(o): return Xform(X.R*o.R,X.R*(o.t) + X.t)
                elif ismat(o):   return Xform(X.R*o,X.t)
                elif islist (o):  return [X*x for x in o]
                elif istuple(o): return tuple([X*x for x in o])
                elif isiter(o):  return (X*x for x in o)
                else:            return o.__rmul__(X)
        def __rmul__(X,o):
                if ismat(o): return Xform(o*X.R,o*X.t)
                raise NotImplementedError
        def __div__(X,o):
                if isxform(o): return X*~o
                return o.__rdiv__(X)
        def __add__(X,v):
                if isvec(v): return Xform( X.R, X.t + X.R*v )
                return v.__radd__(X)
        def __radd__(X,v):
                if isvec(v): return Xform( X.R, X.t + v )
                raise NotImplementedError
        def __sub__(X,v):
                if isvec(v): return Xform( X.R, X.t - X.R*v )
                return v.__rsub__(X)
        def __rsub__(X,v):
                if isvec(v): return Xform( X.R, X.t - v )
                raise NotImplementedError
        def __eq__(self,other): return self.R==other.R and self.t==other.t
        def __str__ (self): return "Xform( %s, %s )" % (str(self.R),str(self.t))
        def __repr__(self): return "Xform( %s, %s )" % (repr(self.R),repr(self.t))
        def __eq__(X,Y):
                assert isxform(Y)
                return X.R == Y.R and X.t == Y.t
        def rotation_axis(X): return X.R.rotation_axis()
        def pretty(self):
                a,r = self.rotation_axis()
                if self.t.length() > EPS: return "Xform( axis=%s, ang=%f, dir=%s, dis=%f )"%(str(a),degrees(r),str(self.t.normalized()),self.t.length())
                else: return "Xform( axis=%s, ang=%f, dir=%s, dis=%f )"%(str(a),degrees(r),str(V0),0)
Ixform = Xform(Imat,V0)
def stub(cen=None, a=None, b=None, c=None):
        if cen is None: cen = a
        if c   is None: cen,a,b,c = cen,cen,a,b
        return Xform().from_four_points(cen,a,b,c)
def enumerate_combinations(list):
    num = len(list)
    ncomb=1
    counter=[]
    for i in range(num):
       ncomb=ncomb*list[i]
       counter.append(0)
    add = 1
    combs=[]
    for j in range(ncomb):
      combs.append(string.join(map(lambda x: str(counter[x]), range(num))))
      for i in range(num):
        counter[i]=counter[i]+add
        if counter[i]==list[ i ]:
           counter[i]=0
        else:
           break
    return(combs)
def Generate_Pdbs(Nres,num_chain,num_to_output,w,R,orientation,helix_phase,delta_z,output_file_name,chain_name, chain_order):
 deg_to_rad= math.pi/180.
# chain  parameters
 ph = 360/num_chain
 phase=[]
 for i in range(num_chain):
     phase.append(i*ph)
 chain_set=['A','B','C','D','E','F','G','H',' I ',' J ',' K','L','M','N','O','P','Q','R','S ',' T','U','V','W','X','Y','Z']
 chain_num=chain_set[0:num_chain]
 R1=2.26
#rise per  residue  d   fixed .  this   constrains   pitch  ( alpha )
 d=1.51
 z1=0.
 z2=0.
 line='ATOM      7  CA  GLY A   2       5.520   2.352   1.361  1.00 20.00     '
 last=line[54:-1]
 atom_num=1
 res_num=1
 Res_id=[]
 CA_list=[]
 alpha=math.asin(R*w*deg_to_rad/d)
 for iter in range(num_to_output):
  CA_list.append([])
  Res_id.append([])
  chain = chain_name[iter]
  orient = orientation[iter ]
  #w1=100-w
  w1=99.2 # decouple w1 and w and fix w1 to 99.2 (3 full turns per 11 residues). Change to 98.2 for TIP-98 or 97.2 for TIP-97 etc.
  if orient == 1:
     res_num=0
  else :
     res_num=Nres+1
  supercoil_phase=phase[iter]+delta_z[iter]*math.tan(alpha)/(R*deg_to_rad)
  for t in range(Nres+2):      ## need two extra residues to  guide   placement  of 1 st  and  last   residue
    a0=(w*(t-1)+supercoil_phase)*deg_to_rad
    if orient == 1:
     a1=(w1*(t-1)+helix_phase[iter])*deg_to_rad   # set ref point for phase to be along  supercoil   radius
    else:
     a1=(w1*t-w1*Nres-helix_phase[iter])*deg_to_rad
    x=R*math.cos(a0) + R1*math.cos(a0) * cos(a1) - R1*cos(alpha)*sin(a0)*sin(a1)
    y=R*math.sin(a0) + R1*sin(a0)*cos(a1) + R1*cos(alpha)*cos(a0)*sin(a1)
    if w==0:
     z=d*t+delta_z[iter]
    else:
     z= R*w*t*deg_to_rad/math.tan(alpha)-R1*sin(alpha)*sin(a1)+delta_z[iter]
    CA_list[iter].append( (res_num,Vec(x,y,z)) )
    Res_id[iter].append(res_num)
    atom_num=atom_num+1
    if orient  == 1:
        res_num=res_num+1
    else:
        res_num=res_num-1
# convert CA trace to  full   backbone  model by  superimposing  on  ideal   template
# by matching 3  consecutive  CA atoms
# set up  ideal   template
 stub_file=map(string.split,open('ideal.pdb','r ') . readlines ())
 atom=[]
 for line in stub_file:
    atom.append( (Vec(float(line[6]),float( line [7]) , float ( line [8]) )))
 ideal_stub=stub(atom[6],atom[1],atom[11])
#now make full  backbone  pdb
 full_pdb=open(output_file_name,'w')
 atom_num=1
 res_num=0
 for counter in range(num_to_output):
  iter=int(chain_order[counter])
  chain=chain_name[iter]
  CA_chain_u=CA_list[iter]
  CA_chain = sorted(CA_chain_u, key = lambda res: res[0])
  for res in range(1,Nres+1):
    res_num=res_num+1
    actual_stub=stub(CA_chain[res][1],CA_chain[res-1][1],CA_chain[res+1][1])
    transform=actual_stub * ~ideal_stub
# N
    coords=transform*atom[5]
    full_pdb.write('ATOM %6d  N   GLY %s %3d    %8.3f%8.3f%8.3f%s\n'%(atom_num,chain,res_num,coords.x,coords.y,coords.z,last))
    atom_num=atom_num+1
# CA   (use actual CA from trace  rather  than  superimposed  one)
    coords=CA_chain[res][1]
    tcoords=transform*atom[6]
#    print  coords , tcoords ,' CA'
    full_pdb.write('ATOM %6d  CA  GLY %s %3d    %8.3f%8.3f%8.3f%s\n'%(atom_num,chain,res_num,coords.x,coords.y,coords.z,last))
    atom_num=atom_num+1
#  NH
    coords=transform*atom[7]
    full_pdb.write('ATOM %6d  H   GLY %s %3d    %8.3f%8.3f%8.3f%s\n'%(atom_num,chain,res_num,coords.x,coords.y,coords.z,last))
    atom_num=atom_num+1
#  C
    coords=transform*atom[8]
    full_pdb.write('ATOM %6d  C   GLY %s %3d    %8.3f%8.3f%8.3f%s\n'%(atom_num,chain,res_num,coords.x,coords.y,coords.z,last))
    atom_num=atom_num+1
# O
    coords=transform*atom[9]
    full_pdb.write('ATOM %6d  O   GLY %s %3d    %8.3f%8.3f%8.3f%s\n'%(atom_num,chain,res_num,coords.x,coords.y,coords.z,last))
    atom_num=atom_num+1
    start_d=Vec.distance(atom[8],atom[6])
    end_d = Vec.distance(transform*atom[8],transform*atom[6])
 return()
def input_params(input):
    output_file_prefix=input[0][0]
    Nres = int(input[1][0])
    w0_begin = float(input[2][0])
    w0_end = float(input[2][1])
    w0_iter=int(input[2][2])
    R0_begin  = float(input[3][0])
    R0_end = float(input[3][1])
    R0_iter = int(input[3][2])
    num_chain=int(input[4][0])
    num_to_output=int(input[5][0])
    orientation=[]
    for i in range(num_to_output):
        orientation.append( int(input[6][i] ))
    phase=[]
    for i in range(num_to_output):   ## restrict phase and delta_z to switch  sign  in sym_mates
        phase.append( (float(input[7+i][0]),float(input[7+i ][1]) ,int(input[7+i ][2]) ))
    z_list=[]
    for i in range(num_to_output):
        l=7+num_to_output+i
        z_list.append( ( float(input[l ][0]) , float (input[ l ][1]) ,int(input[ l ][2])  ))
    chain_name=[]
    for i in range(num_to_output):
        chain_name.append(input[l+1][i])
    chain_order=[]
    for i in range(num_to_output):
        chain_order.append(input[l+2][i])
    print ' ###############    SUPERHELIX PARAMS   ############## \n'
    print ' output file prefix:  %s \n'%(output_file_prefix)
    print ' helix length:  %s \n'%(Nres)
    print ' starting twist: %s  ending twist: %s  number of samples: %s \n'%(w0_begin,w0_end,w0_iter)
    print ' starting R0:   %s   ending R0:  %s   number of smaples:  %s \n'%(R0_begin,R0_end,R0_iter)
    print ' number of chains:  %s \n'%(num_chain)
    print ' number of chains in assymetric unit: %s \n' %(num_to_output)
    print ' ###############  individual helix parameters  ######## \n'
    print ' ORIENTATION CHAIN CHAIN_ORDER   PHASE (start, end, number)   Z OFFSET (start, end, number) \n '
    for i in range(num_to_output):
        print '%s %s %s    %s %s %s              %s %s %s \n'%(orientation[i],chain_name[i],chain_order[i],phase[i][0],phase[i][1],phase[i][2],z_list[i][0],z_list[i][1],z_list [ i ][2])
    # sample p1 evenly , w and R around the  input   values
    number_of_combinations=w0_iter*R0_iter
    for i in range(num_to_output):
        number_of_combinations=number_of_combinations*phase[i][2]*z_list[i][2]
    print  '  NUMBER OF PDBS TO GENERATE:  %s \n'%number_of_combinations
    combinations=[]
    w0_inc = (w0_end - w0_begin)/max(w0_iter-1,1)
    R0_inc = (R0_end - R0_begin)/max(R0_iter-1,1)
    w0=[]
    R0=[]
    ph=[]
    z = []
    for i in range(w0_iter):
        w0.append( w0_begin+w0_inc*i)
    for i in range(R0_iter):
        R0.append( R0_begin+R0_inc*i)
    for j in range(num_to_output):
        p_inc=(phase[j][1] - phase[j][0])/max(phase[j][2]-1,1)
        ph.append([])
        for i in range(phase[j][2]) :
             ph[j].append(phase[j][0]+p_inc*i)
    for j in range(num_to_output):
        z_inc=(z_list[j][1] - z_list[j ][0]) /max(z_list[j][2]-1,1)
        z.append([])
        for i in range(z_list[j ][2]) :
            z[j].append(z_list[j][0]+z_inc*i)
    return(Nres,num_chain,num_to_output,orientation,R0,w0,ph,z,output_file_prefix,chain_name,chain_order)
####################################################
input_file=argv[1]
tag = argv[2]
input =map(string.split,open(input_file,'r') . readlines ())
Nres,num_chain,num_to_output,orientation,R0,w0,ph,z,output_file_prefix,chain_name,chain_order = input_params(input)
items=[]
items.append(len(w0))
items.append(len(R0))
for p in ph:
    items.append(len(p))
for zz in z:
    items.append(len(zz))
combos=enumerate_combinations(items)
for combo in combos:
    id=map(int,string.split(combo))
    helix_phase=[]
    delta_z =[]
    for i in range(num_to_output):
        helix_phase.append(ph[i][id[2+i]])
        delta_z.append(z[i][id[2+num_to_output+i]])
    ## to get close to C2 symmetry for  axis  in x-y plane,  switch   sign  of  phase and  delta_z  of sym mates
    out_file_name='%s_%.2f_%.2f'%(tag,w0[id[0]],R0[id[1]])
    for i in range(num_to_output):
        out_file_name=out_file_name+'_'+'%.2f'%helix_phase[i]
    for i in range(num_to_output):
        out_file_name=out_file_name+'_'+'%.2f'%delta_z[i]
    out_file_name=out_file_name+'.pdb'
    Generate_Pdbs(Nres,num_chain,num_to_output,w0[id[0]],R0[id[1]],orientation,helix_phase,delta_z,out_file_name,chain_name,chain_order)
###################End of script##############################
```

Execution: python2.7 generate_helix.py <design_name> params.dat

***Script S2. Rosetta Design scripts***

Simplified design script and score weight (.wts) adapted from Huang et al. (1) to design helix bundle backbones.

```
design.xml
<ROSETTASCRIPTS>
<SCOREFXNS>
	<ScoreFunction name="soft" weights="/input/soft_rep_trp_ala"/>
	<ScoreFunction name="hard" weights="ref2015">
		<Reweight scoretype="coordinate_constraint" weight="0.5" />
	</ScoreFunction>
	<ScoreFunction name="hard_bb" weights="/input/bb_only">
		<Reweight scoretype="coordinate_constraint" weight="2." />
		<Reweight scoretype="cart_bonded" weight="0.5" />
	</ScoreFunction>
</SCOREFXNS>
<RESIDUE_SELECTORS>
	<Or name="IBP">
<Index resnums="47"/>
<Index resnums="50"/>
<Index resnums="54"/>
<Index resnums="58"/>
<Index resnums="61"/>
<Index resnums="65"/>
<Index resnums="69"/>
<Index resnums="72"/>
<Index resnums="76"/>
<Index resnums="80"/>
<Index resnums="83"/>
<Index resnums="87"/>
</Or>
<Layer name="core" select_core="true" use_sidechain_neighbors="false" ball_radius="2.5" />
<Not name="not_core" selector="core" />
<Layer name="boundary" select_boundary="true" use_sidechain_neighbors="false" ball_radius="2.5" />
<Not name="not_boundary" selector="boundary" />
<Layer name="surface" select_surface="true" use_sidechain_neighbors="false" ball_radius="2.5" />
<Not name="not_surface" selector="surface" />
</RESIDUE_SELECTORS>

<TASKOPERATIONS>
<IncludeCurrent name="current"/>
<LimitAromaChi2 name="arochi" />
<ExtraRotamersGeneric name="ex1_ex2" ex1="1" ex2="1"/>
<OperateOnResidueSubset name="surface_res_allowed" selector="surface"> <RestrictAbsentCanonicalAASRLT aas="DESTNQKRH"/>
    </OperateOnResidueSubset>
    <OperateOnResidueSubset name="notsurface" selector="not_surface">
        <PreventRepackingRLT/>
    </OperateOnResidueSubset>
    <OperateOnResidueSubset name="core_res_allowed" selector="core">
        <RestrictAbsentCanonicalAASRLT aas="VAILMF"/>
    </OperateOnResidueSubset>
    <OperateOnResidueSubset name="notcore" selector="not_core">
        <PreventRepackingRLT/>
    </OperateOnResidueSubset>
    <OperateOnResidueSubset name="boundary_res_allowed" selector="boundary">
        <RestrictAbsentCanonicalAASRLT aas="VAILYDESTNQKRH"/>
    </OperateOnResidueSubset>
    <OperateOnResidueSubset name="notboundary" selector="not_boundary">
        <PreventRepackingRLT/>
    </OperateOnResidueSubset>
    <OperateOnResidueSubset name="restrict_IBP" selector="IBP">
        <RestrictToRepackingRLT/>
    </OperateOnResidueSubset>
	<RestrictAbsentCanonicalAAS name="ala_only" resnum="0" k	eep_aas="A" />
</TASKOPERATIONS>

<FILTERS>
	<PackStat name="packstat" threshold="0.3" confidence="1"/>
</FILTERS>

<MOVERS>
<AddConstraintsToCurrentConformationMover name="add_cst" use_distance_cst="0" max_distance="12" coord_dev="2.5" min_seq_sep="8" />
	<ClearConstraintsMover name="clearconstraints"/>
<PackRotamersMover name="transform_sc" scorefxn="hard" task_operations="ala_only"/>
<PackRotamersMover name="softpack_core" scorefxn="soft" task_operations="core_res_allowed,notcore,current,arochi,restrict_IBP"/>
<PackRotamersMover name="softpack_surface" scorefxn="soft" task_operations="surface_res_allowed,notsurface,current,arochi,restrict_IBP"/>
<PackRotamersMover name="hardpack_surface" scorefxn="hard" task_operations="surface_res_allowed,notsurface,current,arochi,ex1,restrict_IBP"/>
<PackRotamersMover name="hardpack_core" scorefxn="hard" task_operations="core_res_allowed,notcore,current,arochi,ex1_ex2,restrict_IBP"/>
<PackRotamersMover name="softpack_boundary" scorefxn="soft" task_operations="boundary_res_allowed,notboundary,current,arochi,restrict_IBP"/>
<PackRotamersMover name="hardpack_boundary" scorefxn="hard" task_operations="boundary_res_allowed,notboundary,current,arochi,ex1_ex2,restrict_IBP"/>
<MinMover name="hardmin_bb" scorefxn="hard_bb" type="lbfgs_armijo_nonmonotone" tolerance="0.0001" chi="1" bb="1" bondangle="1" bondlength="1" jump="all" cartesian="1"/>
<MinMover name="hardmin_sconly" scorefxn="hard" chi="1" bb="0" bondangle="0" bondlength="0"/>
	<MutateResidue name="50T" target="50" new_res="THR"/>
	<MutateResidue name="60T" target="61" new_res="THR"/>
	<MutateResidue name="70T" target="72" new_res="THR"/>
	<MutateResidue name="80T" target="83" new_res="THR"/>
</MOVERS>
<PROTOCOLS>
	<!--mutate all residues in the pose to alanines-->
	<Add mover="transform_sc"/>
	<!--minimize the backbone with constrains -->
	<Add mover="add_cst"/>
	<Add mover="hardmin_bb"/>
	<Add mover="clearconstraints"/>
	<!--apply threonine mutations-->
	<Add mover="50T"/>
	<Add mover="60T"/>
	<Add mover="70T"/>
	<Add mover="80T"/>
	<!--layer design round 1-->
	<Add mover="softpack_core"/>
	<Add mover="softpack_boundary"/>
	<Add mover="softpack_surface"/>
	<Add mover="hardmin_sconly"/>
	<!--layer design round 2-->
	<Add mover="hardpack_core"/>
	<Add mover="hardpack_boundary"/>
	<Add mover="hardpack_surface"/>
	<!--remove designs with packstat < 0.3-->
	<Add filter="packstat"/>
</PROTOCOLS>
</ROSETTASCRIPTS>
```

soft_rep_trp_ala.wts

ETABLE FA_STANDARD_SOFT

METHOD_WEIGHTS ref 1.0 -0 -0.699 -0.654 1.652 -0.449 0.89 0.526 -0.749 0.226 0.328 -0.877 0.214 -0.834 -0.797 -0.492 -0.743 0.177 2.0 1.077

pro_close 1.0

fa_atr 0.8

fa_rep 0.657

fa_sol 0.648

fa_intra_rep 0

fa_pair 0.452

fa_dun 0.569

ref 1

hbond_lr_bb 0.857

hbond_sr_bb 0.857

hbond_bb_sc 0.857

hbond_sc 0.857

p_aa_pp 0.915

dslf_ss_dst 0.5

dslf_cs_ang 2

dslf_ss_dih 5

dslf_ca_dih 5

bb_only.wts

METHOD_WEIGHTS ref 3.6 -0.000206398 -0.47402 -0.604935 1.15537 -0.310674 0.954298 0.0 -0.442574 0.0 -0.204523 -0.74518 -0.654485 -0.781173 -0.76836 -0.298617 -0.22266 0.025644 2.0 0.928496

fa_rep 0.44

fa_intra_rep 0.004

hbond_sr_bb 1.5

cart_bonded 0.5

rama 0.2

omega 0.5

ref 1

Execution: ./rosetta_scripts.default.linuxgccrelease -parser:protocol design.xml -s input_name.pdb -out:path:all /output_folder -nstruct 1 -overwrite

***Script S3. Rosetta Remodel scripts***

Example of parameters for TIP-99 loop modelling.

remodel.flags

-in:path input_files

-remodel:blueprint input_files/blueprint

-remodel:num_trajectory 1

-nstruct 500

-remodel:quick_and_dirty

-out:path:all output_files

-out:file:scorefile missing_loops.sc

-find_neighbors

-overwrite

-ex1

-ex2

Example of blueprint for TIP-99 loop modelling.

Residues that are remodeled to connect the loops of helices in bold. The two adjacent residues to the loop connection are also remodeled. Only TIP-97 required a 3 residue loop to connect loops. After loop closure in centroid mode the residues are Rosetta designed and neighboring residues to the loops are repacked. In the residue design, all amino acid identities are allowed except for cysteine.

Blueprint

1 D .

2 E .

3 E .

4 E .

5 E .

6 I .

7 K .

8 K .

9 A .

10 E .

11 K .

12 L .

13 V .

14 E .

15 D .

16 L .

17 L .

18 R .

19 E .

20 A .

21 K .

22 K .

23 I .

24 V .

25 E .

26 E .

27 I .

28 L .

29 R .

30 E .

31 L .

32 K .

33 K .

34 A .

35 V .

36 E .

37 E .

38 L .

39 R .

40 E .

41 R .

42 L .

43 E .

**44 K L NOTAA C**

**0 x L NOTAA C**

**0 x L NOTAA C**

**45 D L NOTAA C**

46 E .

47 E .

48 E .

49 A .

50 E .

51 R .

52 I .

53 E .

54 R .

55 E .

56 M .

57 K .

58 R .

59 L .

60 A .

61 E .

62 E .

63 A .

64 K .

65 K .

66 R .

67 M .

68 Q .

69 E .

70 T .

71 A .

72 E .

73 K .

74 L .

75 K .

76 K .

77 E .

78 L .

79 E .

80 R .

81 I .

82 E .

83 K .

84 E .

85 L .

86 R .

87 D .

**88 R L NOTAA C**

**0 x L NOTAA C**

**0 x L NOTAA C**

**89 D L NOTAA C**

90 E .

91 E .

92 E .

93 E .

94 I .

95 E .

96 K .

97 R .

98 A .

99 E .

100 E .

101 A .

102 K .

103 R .

104 R .

105 M .

106 E .

107 E .

108 L .

109 I .

110 R .

111 K .

112 A .

113 K .

114 E .

115 E .

116 M .

117 E .

118 K .

119 L .

120 I .

121 E .

122 K .

123 A .

124 E .

125 R .

126 E .

127 I .

128 K .

129 E .

130 K .

131 N .

132 K .

Execution: ./remodel.default.linuxgccrelease @remodel.flags

***Script S4. Backbone RMSD calculation script***

```
calculate_rmsd.py

##### START OF SCRIPT #####
from pyrosetta import init
from pyrosetta.io import pose_from_pdb
from pyrosetta.rosetta.core.scoring import CA_rmsd
import matplotlib.pyplot as plt

init()
# top_pdb.list contains a list of file names without pdb extention
f = open("top_pdbs.list","r")
out=open("rmsd_pdbs.dat","w+")

list_pdbs = []
list_rmsd = []
for line in f:
		native = line.splitlines()[0]+".pdb"
		relaxed = line.splitlines()[0]+"_0001.pdb"
pose_native = pose_from_pdb("/input/"+native)
pose_relaxed = pose_from_pdb(“/output/"+relaxed)
		rmsd = CA_rmsd(pose_native,pose_relaxed)
		list_pdbs.append(native)
		list_rmsd.append(rmsd)
		out.write(native+"\t"+str(rmsd)+"\n") #write to file
f.close()
out.close()
##### END OF SCRIPT #####
```

**Script S6. Rosetta *Abinitio* structure prediction**

3-mer and 9-mer were generated via the Robetta web server (12). The input.fasta contains the designed amino acid sequence in fasta format. Typically >3000 AbinitioRelax simulations are perfomed to observe a folding funnel with the following flags:

abinitio.flags

-in:file:native input_files/design.pdb

-in:file:fasta input_files/input.fasta

-in:file:frag3 input_files/frags.200.3mers

-in:file:frag9 input_files/frags.200.9mers

-nstruct 50

-abinitio:relax

-abinitio::increase_cycles 10

-abinitio::rg_reweight 0.5

-abinitio::rsd_wt_helix 0.5

-abinitio::rsd_wt_loop 0.5

-relax::fast

-out:file:silent output_files/fold_silent_${SLURM_ARRAY_TASK_ID}.out

-overwrite

Execution: ./AbinitioRelax.default.linuxgccrelease @abinitio.flags
